# Interpretable machine learning reveals a diverse arsenal of anti-defenses in bacterial viruses

**DOI:** 10.1101/2024.06.14.598830

**Authors:** Anna Lopatina, Mariusz Ferenc, Nina Bartlau, Michael Wolfram, Kerrin Steensen, Sebastiano Muscia, Fatima Hussain, Madeleine Schnurer, Leila Afjehi-Sadat, Shaul Pollak, Martin Polz

## Abstract

Antagonistic interactions with viruses are an important driver of the ecology and evolution of bacteria, and associating genetic signatures to these interactions is of fundamental importance to predict viral infection success. Recent studies have highlighted that bacteria possess a large, rapidly changing arsenal of defense genes and that viruses can neutralize at least some of these genes with matching anti-defenses. However, a broadly applicable approach for discovering the genetic underpinnings of such interactions is missing since typically used methods such as comparative genomics are limited by the rampant horizontal gene transfer and poor annotation of viral and bacterial genes. Here we show that genes that allow the viruses to overcome bacterial defenses can be systematically identified using an interpretable machine-learning approach even when using diverse bacteria-virus infection data. To verify the predictions, we experimentally characterized eight previously unknown anti-defense proteins in viruses specific for *Vibrio* bacteria and showed that they counteract a wide range of bacterial immune systems, including AbiH, AbiU, Septu, DRT, CBASS, and Retron. The power of our computational approach is highlighted by the identification of anti-defense proteins that inhibit non-homologous defense systems, which we verify for Retron and AbiH. We suggest that the computational prediction based on experimental interactions offers a promising avenue to unravel the genetic mechanisms of co-evolution between bacteria and their viruses.

## Introduction

While viruses are thought to exert control over bacterial populations^1,2^, the discovery of a diverse arsenal of defense genes within bacterial genomes questions how such control may be achieved in the wild. To date, over 130 defense systems have been identified within bacterial isolates, sometimes containing more than a dozen different defense systems each^3–5^. Their combined action has been termed pan-immunity because they cluster in the flexible genome where they turn over rapidly^3,6^. That such defense of bacteria is effective was confirmed through large virus-bacteria interaction matrices, revealing sparse lytic interactions^3,7–10^. However, these matrices also suggested that closely related viruses can overcome host defenses by various mechanisms such as point mutations or anti-defense genes^11–13^. Indeed, comparative genomics of largely identical viruses often demonstrates synteny interrupted by regions of high turnover of diverse genes that are predominantly short and lack annotation^14^. This feature of closely related viruses was recently utilized to discover anti-defense genes against several types of defenses, which were named anti-CBASS^15–17^, anti-Thoeris^18,19^, anti-DSR2^20^, anti-Gabija^21^ and anti-Pycsar^16^. However, more generally, associating genetic features of more divergent viruses and their hosts is hampered by several problems that render comparative genomics and even genome-wide association studies ineffective, including the co-existence of multiple defenses in the same bacterial genome, their high evolutionary turnover by gene transfer and loss, as well as the general lack of anti-defense gene annotation. Because large cross-infection matrices have recently become a common tool to study bacteria-virus interactions, we asked to what extent computational approaches hold the power to reveal the genetic underpinnings of the arms race of defense and anti-defense from such cross-infection matrices.

Our test case for computational prediction of defense and anti-defense gene interactions is the Nahant cross-infection matrix, which consists of 65,232 experimentally determined interactions of co-occurring *Vibrio* bacteria and their viruses isolated from a coastal ocean environment. Because the matrix contains a range of closely and distantly related viruses, we first explored the extent to which comparative genomics can discover defense and anti-defense gene pairs (Methods). We therefore first annotated the diversity of bacterial defense genes in the Nahant collection of 758 fully sequenced *Vibrio* genomes. Using DefenseFinder^4^, we detect 50 defense systems in 5,623 operons, providing a good representation of the −60 defense systems that were available in version 1.0.8 of the database (Supplementary Data 1). Our analysis shows that although cases of putative anti-defense genes in viral genomes can be identified, systematically pairing them with known defense genes in bacterial genomes is impossible because of the co-occurrence of multiple defenses in the bacterial isolates (Extended Data Fig. 1), a problem that could only be solved by extensive trial and error cloning. We reasoned that because there are experimentally verified infection data that can be combined with bioinformatic identification of defense genes, their cognate viral anti-defense genes can be efficiently predicted by a supervised machine learning approach. Our specific hypothesis is that if infection outcomes are predictable from the genomic information of each bacterial-viral pair, an interpretable machine learning approach will allow identification of the viral genes that overcome host defenses. To develop and train our method, we augmented the Nahant cross-infection matrix with genome-resolved infection outcomes from the literature and with known defense and anti-defense pairs for validation (Methods). By subsequent experimental verification, we show that the method allows accurate pairing of defenses and anti-defenses, including the surprising finding that commonly a single anti-defense can act against multiple, non-homologous bacterial defense types.

## Results

### Computational predictions

At the core of our computational approach is a convolutional neural network (CNN), which is trained to predict infection outcomes and based on known bacterial defenses, allows us to identify novel anti-defenses using interpretable machine-learning. The input data for each bacteria-virus pair required by the CNN is the infection outcome label and a 2-column array containing bacterial defense proteins and all viral proteins (Fig. 1A). Because CNNs require numeric data as input, we use a language model to represent proteins numerically in these input columns. Due to its proven ability to capture structural, functional, and evolutionary information in its numeric representation of proteins (embeddings)^22–24^, we use a bidirectional encoder representation transformer (BERT) architecture^25^. The ability of our embeddings to capture functional information is evident by the clustering of proteins belonging to the same defense system type in our embedding space (Fig. 1B). By concatenating embeddings around each focal protein (Fig. 1C), we augment embeddings by including two evolutionary features - the clustering of functionally related proteins on the chromosome and the tendency of homologous sequences that have evolved different activities to occur in different genomic backgrounds. Our CNN achieves high performance in predicting infection outcome labels solely based on information on bacterial defenses and virus proteins across our training and validation datasets (average 5-fold cross-validation accuracy = 0.942). We then learn which bacterial defense and (unknown) viral proteins contributed most to the infection prediction in the CNN by performing a Shapley value-based analysis^26^ (Methods), which allows us to rank putative anti-defense proteins according to a score based on normalized Shapley values (ranging from 0-100) (Fig. 1D). We report putative anti-defenses with a score >50 since results for our validation dataset provided the score distribution for 9 previously verified defense and anti-defense pairs from 6 defense system types and showed that 50 was the lowest score for a true positive (Supplementary Table 1).

**Figure. 1.**
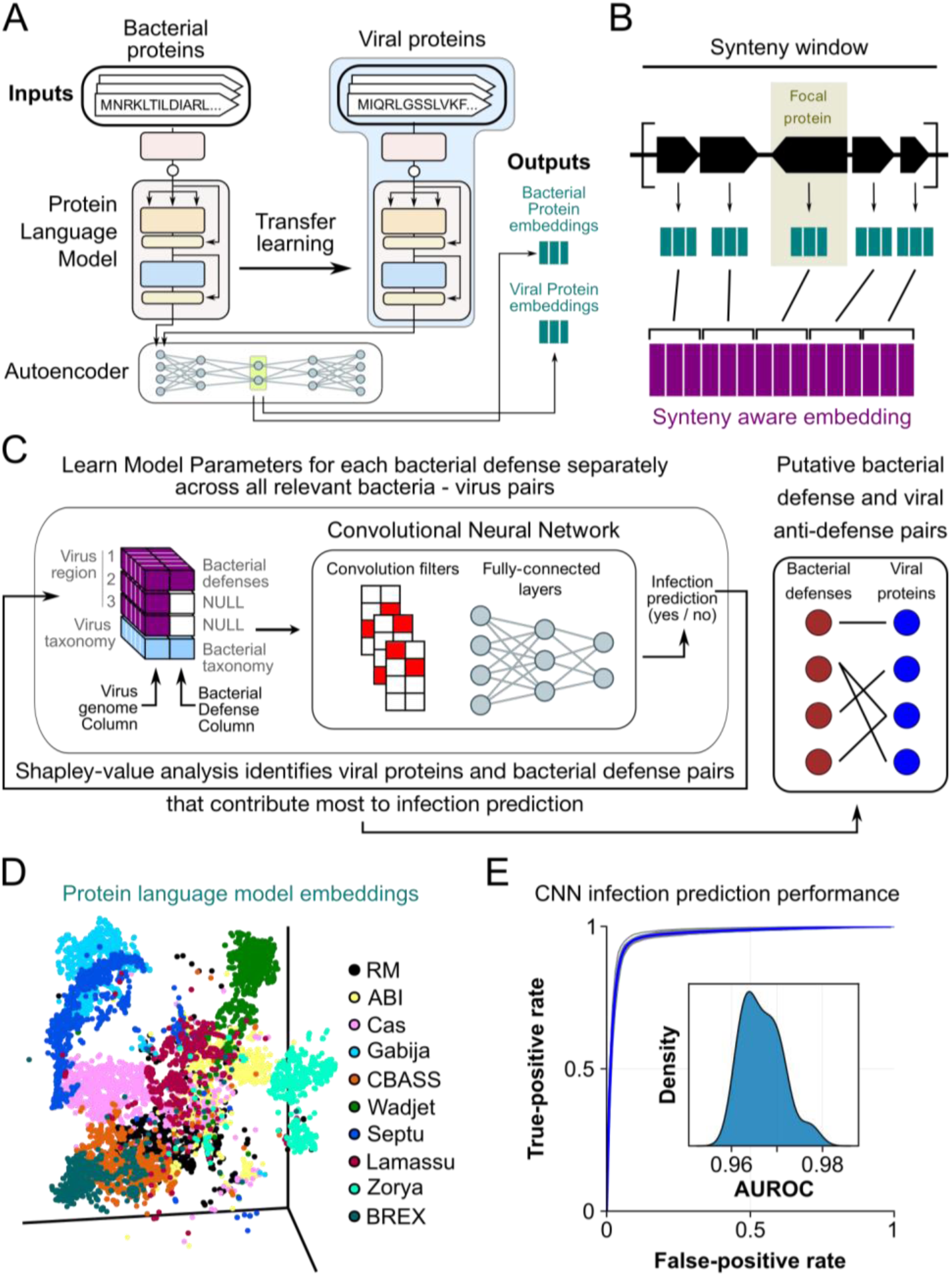
Machine learning approach to identify anti-defense genes in bacterial viruses. **a,** A BERT protein language model with ∼4.2*10^8^ parameters trained on >10^8^ bacterial proteins converts protein sequences from string representations (top) to 1024 dimensional numeric embeddings (represented as arrows going into the autoencoder block). The network is re-trained on viral proteomes using transfer learning approaches to produce viral protein embeddings. Both bacterial and viral protein embeddings are then passed through an autoencoder to produce the final 128-dimensional embeddings (Outputs) used in downstream tasks. **b,** Final embedding vectors take local genomic context into account by concatenating 19 embeddings on each side of the focal protein according to their order on the chromosome (methods). **c,** Viral and bacterial genomes are arranged into the columns of a 4×2×2434 tensor (left). The first 3 rows of the viral column each represent an equal partition of the viral genome, while the 4^th^ row represents viral taxonomy (methods). The bacterial column contains a single synteny-aware vector containing all defenses in the order they appear on the genome, two blank vectors, and a taxonomy vector in the fourth row. A convolutional neural network is then trained to predict infection outcome labels from all coupled viral-bacterial inputs. We identify features in each input tensor that contribute most to infection prediction using a Shapley value analysis, only retaining cases where a viral protein received a normalized score >50 as putative anti-defenses. Inhibitory links between specific defense and anti-defense pairs are inferred as cases where both viral and bacterial features received high scores, allowing multiple connections per protein as shown in the network on the right. **d,** 3D visualization of protein embeddings of examples of bacterial defense systems. Each point represents a single protein, and colors depict defense system membership as predicted by DefenseFinder v1.0.8. Axes represent latent dimensions of the innermost layer of an autoencoder trained to reconstruct defense system membership from input embeddings. **e,** CNN performance ROC curves. The main plot shows the mean ROC curve for all CNNs trained on different focal bacterial defenses in blue with individual models in gray. The inset shows the distribution of AUROCs for the different models with a mean 0.967 and standard deviation 0.004.

We prioritized 19 defense system types for the prediction of novel anti-defenses, because they showcase a variety of protein domains and mechanisms of action (see Methods for justification, and Supplementary Table 2 for the list of systems). These include systems that are highly prevalent in bacterial genomes as well as less common systems with no known anti-defenses as test cases. We retrieved 416 unique pairs of defense and anti-defense genes (i.e., occurring in different bacterial and viral genomes) for these 19 defense systems (summarized in Supplementary Table 3, more detailed information in Supplementary Data 2). Out of the 416 unique pairs for the vibrios and their viruses, 91 viral proteins had a unique amino acid sequence, of which 70 were so divergent (<70% amino acid similarity) that they likely represent different functions and thus putative novel anti-defenses (Supplementary Data 2). These viral proteins are generally short (median length = 79 a.a) and in proximity to other small, unannotated genes, but with no consistent location in the viral genomes (Extended Data Fig. 2). We next attempted to validate a subset of the predictions, including the perhaps most surprising outcome that a single anti-defense protein can inhibit multiple non-homologous defenses.

### Experimental verification of defense-anti-defense interactions

We chose 10 defense system types for experimental validation including those for which (1) no known anti-defenses exist (dGTPase, Septu, DRT, AbiH, Zorya, Avs, AbiU, and Lamassu), and (2) some anti-defenses have been characterized but our model predicts novel anti-defenses (CBASS and Retron^15,16,27^) (Supplementary Table 4). To overcome potential problems of heterologous expression in non-native hosts, we created a cloning host from the Nahant collection isolate (*V. cyclitrophicus* 10N.239.312.A01) by deleting all mobile genetic elements containing putative defenses, allowing expression of defense and anti-defense genes from closely related, co-occurring bacteria (Extended Data Fig. 3A). We refer to this engineered strain as the “superhost” and verified that it displays more uniform sensitivity towards a diverse set of viruses due to a lack of defenses (Extended Data Fig. 3B). Using the superhost strain we isolated 40 additional viruses using the archived viral concentrates from the same environmental samples (Methods) based on the rationale that the deletion of defenses should allow more unbiased isolation of viruses. Overall, we find that the combination of deletion of defenses and pairing of native hosts and viruses resulted in a higher proportion of defenses for which activity could be verified than was observed in the past in heterologous expression^28^. Of 18 cloned defense operons, 16 showed defense phenotypes with >=1 order of magnitude protection against at least one virus, representing 9 out of 10 prioritized systems (Fig. 2, Extended Data Fig. 4). While most defenses were active in the superhost, Zorya type I, DRT type I, dGTPase, and three subtypes of Septu displayed weak or no defense. We were, however, able to verify the defensive function for Zorya type I, DRT type I in a different host-virus system - a *V. lentus* strain which lacks three mobile genetic elements containing several defenses each (*V. lentus Δ3*), and 12 co-isolated viruses (Methods)^3^. Having experimentally verified defense functions, we sought to evaluate the model predictions for anti-defenses by comparing virus infection success in three strains expressing (1) defense genes, (2) defense and anti-defense genes, or (3) GFP (no defense control).

**Figure. 2.**
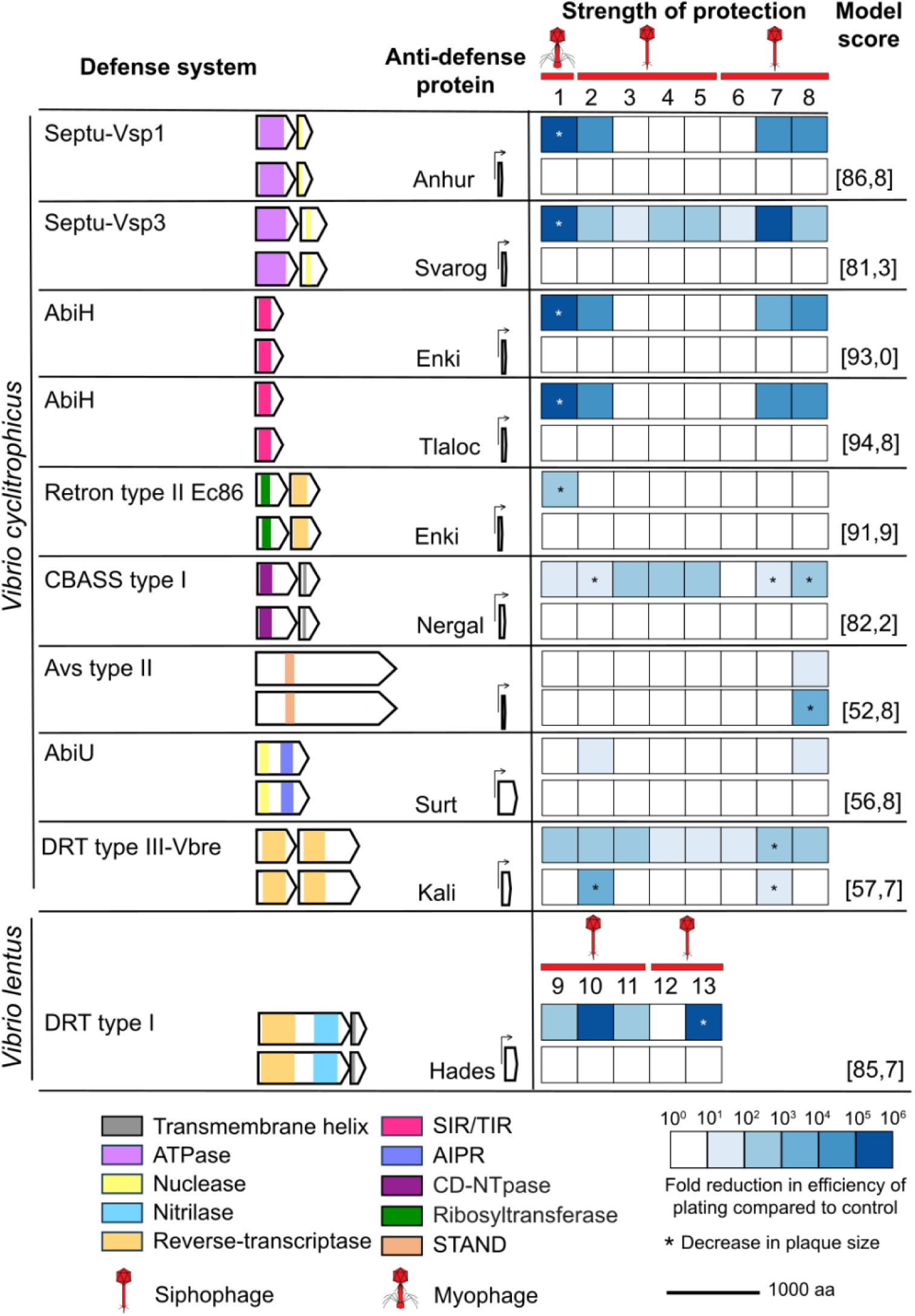
Verification of inhibitory effect of putative anti-defense proteins. The heatmap quantifies host lysis upon challenge with diverse viruses when hosts express a defense system alone or a paired defense system and anti-defense protein from a plasmid. Each column represents a group of viruses (52 total) clustered by genome similarity (Supplementary Table 16) and the values correspond to the average fold change in efficiency of plating (measured in serial dilution plaque assays) compared to an unprotected host (GFP expressing plasmid) for each group of viruses. Asterisks (*) indicate significant reduction in plaque size. Bacterial hosts were *Vibrio cyclitrophicus* superhost or *V. lentus* Δ3, and defenses and anti-defenses were cloned under their native and an inducible PlacI promoter, respectively. Arrow gene diagrams with predicted protein domains in color are drawn to scale.

Experimental verification shows that the model accurately predicts the action of virus encoded anti-defenses (Fig. 2, Extended Data Fig. 5, Supplementary Fig. 1). We successfully cloned 10 of the putative anti-defenses for the 16 bacterial defense operons for which we observed a reduction in virus infection. Of these 10, all seven anti-defenses with high scores in our model (>80) could completely neutralize the action of the defense for at least one virus as indicated by infection levels comparable to the control. Of the two anti-defenses with lower model scores (50ies), one putative anti-defense eliminated the effect of its cognate defense system; one had a protective effect for 6 out of 8 clusters of phages), while the third did not inhibit the defense system. In combination, these results suggest that a score cutoff of 80 yields highly accurate predictions, while lower scores may be less reliable (Supplementary Table 5). As an experimental control, we also cloned one pair of defense and anti-defense not predicted by the model and did not observe a phenotype. Moreover, mass spectrometry showed that defense proteins were expressed independent of the co-expression of the anti-defense from the same plasmid (Supplementary Data 3), providing additional confidence for the validity of the predictions.

Because we noted that a surprising and relatively common feature of the model predictions was that a single anti-defense protein could have multiple high-scoring defense systems that show no apparent homology (Extended Data Fig. 6), we highlight the experimental confirmation of one case where the defenses AbiH and Retron share Enki as a common inhibitor. The 52-amino acid viral protein, predicted to contain a zinc ribbon domain and encoded by the tailless phage 1.080.O (*Autolykiviridae*), can overcome 5.7 orders of magnitude protection conferred by AbiH and 3.2 orders of magnitude protection by Retron. The AbiH defense system is a one-gene system^29^ with a SIR2-like domain previously described as being involved in bacterial immunity and found in multiple different defense systems such as Defense-associated sirtuins (DSRs)^20^, prokaryotic argonautes (pAgo)^30,31^, Thoeris^32^, Avs and additional systems^33^. The Retron system contains a ribosyltransferase effector gene, reverse-transcriptase gene, and msd-msr encoding sequence, and this type of Retron was previously described as a TA system active against phage infection^27^. Although mechanistically unrelated, both defense systems have been reported to cause abortive infection. It is, therefore, possible that this commonality may provide an attack point for this anti-defense protein. A further common occurrence in the model predictions is that unrelated anti-defenses exhibit redundant inhibition of the same defense system. We confirmed such a case for the same AbiH system where the first anti-defense protein (Tlaloc) is a 59-amino acid protein encoded by phage 1.056.O of the *Siphoviridae* family, while the second protein (Enki) is the same 52-amino acid protein that can also inhibit the Retron system. These examples thus show that bacterial viruses have evolved multiple solutions to inhibiting the same defense system, either by having multiple targets or proteins of different evolutionary origins.

Because our predictions frequently did not yield a clear one-to-one relationship for defenses and anti-defenses, we posit that the current practice of naming anti-defenses by using the name of a defense with the addition of “anti” (e.g., Dad as for DRT anti-defense) may lead to confusion. Instead, we follow the previous naming of defense pathways according to gods of protection from different religious traditions and propose names of gods who wreak some form of havoc since the anti-defense genes ultimately enable viruses to destroy their host. Accordingly, we name the validated anti-defenses as Anhur and Svarog against Septu, Kali and Hades against DRT, Tlaloc against AbiH, Enki against AbiH and Retron, Surt against AbiU, Nergal against CBASS (see Supplementary Table 6 for a glossary of the names). However, we note that there are many additional high-scoring defense system-anti-defense protein pairs that can be validated in the future, and since many gods are vengeful, we believe that ample names for anti-defenses will be available.

## Discussion

Our method predicts a vast arsenal of viral anti-defenses, many of which may are generalists, neutralizing several unrelated bacterial defenses. Moreover, specific defense systems can have multiple anti-defenses that appear unrelated, together revealing the complexity of interactions of viruses and bacteria and challenging the one-to-one paradigm of defense and anti-defense interactions. How anti-defenses achieve broad action will be an important avenue of future research as it also pertains to engineering viruses that can overcome a maximum of defenses for phage therapy. While bacteria can expand their genome and harbor many different defenses, viral genomes are highly constrained in length because they need to be packaged into capsids, a problem that is solved by anti-defenses being small and possibly broadly acting proteins.

Because the anti-defenses tend to be in the same overall syntenic location in related viruses, they can be exchanged through homologous recombination of the flanking regions during co-infections^8^, thus ensuring that a population of viruses has pan-immunity, the ability to overcome rapidly changing defenses in their hosts.

Viral predation is an important factor in structuring ecosystems and is gaining increasing prominence as a therapeutic approach to circumvent or supplement the use of antibiotics in combating pathogens. However, the predictability of viral interactions with bacteria is still limited by the lack of a full understanding of the genetic underpinnings of predation success. We show that relatively simple data of positive or negative infection outcomes combined with genomics can be exploited to precisely pinpoint the genes that allow viruses to overcome bacterial defenses, alleviating the need for laborious genetic screens. Our method could be extended to predict viral receptors and synergistic effects of multiple defenses in the same genome by including data on the viral titer necessary to overcome defenses. In this way, the interaction of viruses and bacteria could be highly predictable from sequence data alone, greatly increasing our understanding of environmental dynamics or enabling the more rational design of engineered phages or viral cocktails for phage therapy. In fact, our method can be extended to other systems where high diversity in interactions is due to rapid evolutionary changes, such as in plants where immunity proteins such as Leucine-Rich Repeat domain-containing proteins (LRR) and Pattern Recognition Receptors (PRRs) directly interact with pathogen proteins which are hyper-diverse and have high evolutionary turnover rates. Overall, the vast scope of bacterial diversity usually poses a challenge for functional studies, but it is this diversity that allows interpretable machine learning approaches to make highly accurate predictions, paving the way to associate variation to function more broadly.

## Methods

### Computational analysis

#### Viral and bacterial genome used in the model

A full list of bacterial and viral genomes used for machine-learning tasks is provided in Supplementary Data 4. It consists of publicly available bacterial (n = 4295, Supplementary Data 4) and viral (n = 1949, Supplementary Data 4) genomic nucleotide sequences that were downloaded from NCBI in January 2022. These were supplemented with the 772 *Vibrio* genomes of Nahant collection which were fully sequenced and available in Polz lab on the date of March 2022. Viral genomes were supplemented with 143 genomes of *V.lentus* phages isolated in the lab (see isolation of additional Vibrio bacteria and phages).

#### Viral and bacterial genome annotation

For comparative genomics and machine-learning tasks the pre-existing annotations for all genomes were discarded and open reading frames were called using Bakta v1.9.2^34^ for bacteria and PHANOTATE v1.6.4^35^ for viruses. We used DefenseFinder^4^ v1.0.8 with default parameters to search for prokaryotic antiviral systems in 758 *Vibrio* genomes of Nahant collection which were fully sequenced (a list of all genomes used in this analysis and corresponding taxonomic information is available in Supplementary Data 4).

For the viruses used to challenge the superhost, annotations were done using VirClust^36^ with standard parameters and the following databases: Efam^37^, Efam_XC^37^, PHROG^38^, VOGDB^39^. pVOGs^40^, InterProScan^41^. A consensus annotation was manually curated. Prediction of tRNAs and tmRNAs were done with tRNAscan-SE 2.0^42^ and Aragorn^43^. For the phages isolated on *V. lentus* strains annotations were done using the RAST Server^44^.

To classify the newly isolated phages, relatives were searched by running blastp with standard parameters^45^ against the INPHARED database containing mostly complete genomes^46^ (March 2024). The closest relatives were predicted and filtered by i) evalue < 0.00001; ii) bitscrore > 50; iii) at least 5% of proteins needs to be shared with the query; iv) genome length is at least 50% of smallest query and most 150% of largest query genome. A tree of the subset of relatives with the new isolates was calculated with ViPTree^47^ and only relatives, which shared a branch with the query were chosen. Genus (70% nucleotide similarity) and species (95% nucleotide similarity) clusters were determined by using VIRIDIC^48^. In addition, families were defined by using protein clusters defined with VirClust^36^ with standard parameters (Supplementary Data 5). Gene composition shown in genome comparison plots were created using clinker (version v0.0.28) (Supplementary Fig. 2).

#### Comparative genomics for prediction of viral anti-defense and bacterial defense pairs

Because the Nahant cross-infection matrix contains clusters of closely related viruses, which differ in their ability to lyse the same host or closely related host groups^8^, we explored whether comparative genomics can pinpoint viral anti-defense specific for bacterial defense genes. We therefore first identified groups of closely related viruses and hosts in the matrix using a phylogenetic clustering approach. We calculated average nucleotide identity (ANI) matrices using the command “skani triangulate -s 10 -m 10 -c 10 -E” from Skani v0.2.1. From the underlying distances (defined as 1-ANI*AF), we created a network where nodes are genomes and edges represent distances ≤0.05. To find closely related viral and bacterial groups with high synteny, we clustered the network of vibrios and viruses separately using the Leiden algorithm^49^. To understand the sensitivity of our approach to protein clustering granularity (family, superfamily, etc), all viral and bacterial proteins (see ‘viral and bacterial genome annotation’) were clustered at 0, 30, 50, 70, 80, 90, 95, 99, and 99.9% identity thresholds using Diamond v2.1.9^50^ with the commands “diamond deepclust” and “diamond recluster” using the “--member-cover 60 --masking none --motif-masking 0 --evalue 10” flags and an appropriate “--approx-id” value.

To identify putative anti-defense genes in virus genomes, we next analyzed each closely related virus group by searching for cases where a single bacterial host is infected by some but not all viruses within the group. Based on the assumption that this variability in infection success is due to bacterial gene and viral gene interactions, we define putative anti-defense genes as those present in members of a cluster of closely related viruses that are able to infect the host but absent in those that cannot. Each of these putative anti-defense is paired with bacterial defense systems encoded in the bacterial host genome. If the host itself is a member of a cluster of closely related vibrios in the matrix, we only consider defenses that are found in all members of the *Vibrio* group that are not infected by the closely related viruses. We ran this analysis for every protein homology threshold listed above, resulting in the networks shown in Extended Data Fig. 1.

#### Predicting defense and anti-defense pairs using interpretable machine learning

Here we provide an in-depth description of each step going from raw protein sequences to putative bacterial defense viral anti-defense pairs: (1) Converting proteins to numeric representations using protein-language models, (2) Learning to predict infection outcomes from genomes using CNNs, and (3) Identifying defense and anti-defense pairs using interpretable machine-learning.

Proteins (n=2.55*10^8^ from 5068 bacterial genomes representing 996 genera, and 2381 viral genomes infecting 137 bacterial genera) were converted from string representations to k-mer count vectors, and these were used as input for a BERT language model^25^ as implemented in the PyTorch^51^ library. The model includes 30 BERT blocks, where each block is a composition of attention, normalization, and fully-connected layers (including skip connection and normalization mechanisms). Because bacterial and phage proteins occupied very different areas of embedding space, we chose to learn separate embedding for bacterial and phage proteins each producing 1024 dimensional embeddings. We first trained a BERT model on bacterial proteins, and then used transfer learning techniques to obtain the viral protein embeddings. To reduce computational workload in downstream tasks, we then compressed the embeddings of each model to a final size of 128 features using an autoencoder with 5 fully-connected hidden layers.

We enhance standard embedding techniques to maximally differentiate proteins of different functions using 2 steps: first, by considering the order of genes on the chromosome (synteny), and second, by removing the effect of phylogeny on embeddings. Synteny, where protein families appear in a conserved order on the chromosome, adds information on protein function because of two evolutionary features - the clustering on the chromosome of functionally related proteins and the tendency of homologous sequences with divergent activities to occur in different genomic backgrounds. To incorporate synteny information into embeddings, the BERT output embeddings were concatenated in a small genomic window around each focal protein (windows size = 19 proteins). For bacteria, we only consider the synteny of defense systems identified by DefenseFinder on the chromosome, while for viruses we encode all proteins. To partially remove the effect of phylogeny on the position of proteins in our embedding space, we append a numeric vector that represents taxonomy to protein embeddings in all downstream learning tasks. We first transform each genome nucleotide sequence into a “chaos game representation” (CGR)^52^, which is well-suited for machine-learning applications because of their small and fixed size, and ability to capture sequence patterns at different resolutions. A CGR is created by assigning each of the 4 nucleotides to a corner of a square and mapping the sequence iteratively to this space by using the following procedure: starting at the center, we read the sequence nucleotide by nucleotide, each time placing a dot half-way from the current position to the edge corresponding to the read nucleotide. As shown in Supplementary Fig. 3, this construction creates quadrants and sub-quadrants of a CGR representing k-mers of different lengths. We then trained a convolutional neural network to predict NCBI taxonomy from CGRs (genus for bacteria with weighted-accuracy = 0.99, and family for phages with weighted-accuracy = 0.86). The final representation, which contains synteny-aware embeddings and taxonomy, was used in all downstream learning tasks.

To learn how interactions between viral and bacterial genes determine infection outcome we used a convolutional neural network, as implemented in Tensorflow 2.6.0, where the input is a (4 x 2 x 2432) tensor array, and the output is the infection outcome. We re-train the network for each focal bacterial defense system (see below). Our network has a 16-channel convolutional layer of dimension 3×2×196, a global max-pooling layer, 1 dense layer and a sigmoid output layer. In the input array, each entry is a set of coordinates in the protein embedding space, where rows represent different proteins or taxonomy embeddings, and columns represent the different organisms (virus or bacteria). In the bacterial column, the first row represents the embedding of the different known defense systems found in the strain’s genome centered around the focal system. The 2nd and 3rd rows are assigned blank entries, and the fourth row is populated with the bacterial taxonomic embedding vector. For the virus column, because we do not have a-priori knowledge of genes that may be important for determining infection outcome, we generate 3 embeddings by splitting the virus genome into 3 equal parts and concatenating the embeddings of all proteins in each part. Like the bacterial column, the 4th viral row is also populated by the taxonomy vector of the virus.

We used a Shapley-value analysis^26^ to gain insight into the protein pairs that influence prediction outcome. The Shapley-value is computed for each feature in each input array and represents how much a specific feature, in the context of an input array, contributes to the prediction made by the network. Because the absolute magnitude of the Shapley value is uninformative, we first normalize the Shapley values to the range 0-1, and then order bacteria-virus pairs by the maximal Shapley-value any viral feature achieves in that pair. We then look for input instances where both viral and bacterial features show high (>0.5) normalized Shapley scores, pairing these viral and bacterial proteins as putative anti-defense and defense pairs while assigning a score to the pairing equal to the Shapley value of the viral protein.

### Statistics

Statistical analyses and graphical representations were performed in R^53^ (v.4.2.1) using base R statistical functions and ggplot2^69^ (v3.5.1). To assess differences between two groups, the student two sided t-test was performed.

### Experimental analysis

#### Bacterial growth media

For standard propagation, *Vibrio* isolates were grown at 25°C with 250 rpm shaking in BD Difco 2216 marine broth or in TSB1 medium (Difco Tryptic Soy Broth, 1.5% BD Difco Bacto Agar 214010, amended with 5 g NaCl) for conjugation protocol. *Escherichia coli* strains were grown in BD Difco Miller Luria-Bertani broth (LB) at 37°C with 250 rpm shaking and supplemented for auxotroph strain *E. coli* pi3813 with thymidine (0.3 mM), and for strains *E. coli* β3914 with diaminopimelic acid (dapA) (0.3 mM). Antibiotics were used at the following concentrations: erythromycin (Erm) 200 μg mL-1, kanamycin (Km) 50 μg mL-1 and chloramphenicol (Cm) at 5 or 25 μg mL-1 for *Vibrio* and *E. coli*, respectively.

#### Bacteria and viruses of the Nahant collection

The Nahant cross-infection matrix used in this study consists of bacteria of the genus *Vibrio* and their viruses, assembled in 2010 in the context of a daily time series stretching over 93 consecutive days^8,54,55^. Briefly, bacteria and co-existing viruses were isolated on 3 days (ordinal days 222, 261, and 286) within the 93-day period by direct plating so that each bacterial or viral isolate originates from a colony or plaque-forming unit, respectively, reflecting the abundance of predominant genotypes at the time of sampling. All 259 bacterial isolates for which at least one virus was obtained were subsequently used in an all-by-all cross-infection assay using all 248 independent viral isolates. All bacteria and viruses were genome sequenced and are referred to as the original Nahant collection. We supplemented 259 bacterial isolates with 502 *Vibrio* strains reported in the study^56^, which were fully genome sequenced using Illumina HiSeq (Biomicrocenter, Massachusetts Institute of Technology) with the protocols described earlier^56^. All Vibrio strains are listed in Supplementary Data 4.

#### Isolation of additional Vibrio bacteria and viruses

For this study, we supplemented the original Nahant collection with bacterial and viral isolates. Bacterial isolates were obtained from archived samples for one additional day (ordinal day 239) because tag-sequencing of the rRNA gene indicated that vibrios were undergoing a population expansion around this day^55^. Archived samples consisted of 30% glycerol-BD Difco 2216 marine broth stocks stored at −80°C that contained 0.2 µm filters through which ∼1L of 0.63 µm prefiltered seawater had been passed. Before freezing, the filters in the glycerol stock were shaken extensively to detach cells, and a small portion of the frozen glycerol solution was taken from the tubes and plated on Vibrio-selective MTCBS plates (BD Difco TCBS Agar 265020, supplemented with with 10 g NaCl per liter to 2% final w/v). Colonies were picked, purified by restreaking three times alternatingly on BD Difco 2216 marine broth and MTCBS, and regrown in liquid media (BD Difco 2216 marine broth). Their identity was assessed by sequencing the *hsp60* protein-coding gene, which has served as a marker gene for identifying vibrios, using the primer h279 and h280^57^. Confirmed *Vibrio* isolates were subsequently genome sequenced with a combination of short and long-read sequencing using Illumina HiSeq (Biomicrocenter, Massachusetts Institute of Technology) and Nanopore (Joint Microbiome Facility, University of Vienna) sequencing, respectively (see below “superhost genome sequencing”). Of these isolates, *Vibrio cyclitrophicus* strain 10N.239.312.A01 was chosen for deletion of mobile genetic elements harboring genes predicted to have virus defense functions [see ‘engineering of a defense-less cloning host (superhost)’]. We refer to this strain as “superhost” as it should allow viral isolation and testing of infection efficiency unbiased by genes that can inactivate infecting viruses.

Given the above rationale, we used the superhost as bait for isolation of 40 additional viruses obtained from Julian days 236, 237, 238, 239, 262, 282, and 288 from the Nahant Time Series (Supplementary Table 7). Moreover, we used five isolates of *V. lentus* (strains 10N.286.52.E12, 10N.286.54.F7, 10N.286.54.A6, 10N.261.45.B10, 9CS106) to obtain viruses from Julian days 228, 234, 239-243, 243, 246, 248, 253-263, 266, 267, 269-272, 274-277, 279, 281-283, 285-287, 289-293, 295, 297 and then used these viruses to construct a cross-infection matrix of 40 viruses and 25 strains to supplement the existing Nahant matrix for our computational prediction of anti-defense genes (Supplementary Fig. 4).

All viral isolations from archived concentrates (prepared as previously described^8^) were done by either directly picking viral plaques from agar overlays or by first performing a liquid enrichment followed by plate-based purification. For the first method, 3 ml molten 0.4% top agar (BD Difco 2216 marine broth) at 45°C was mixed with 100 µl of bacterial overnight culture (grown at 25°C and 250 rpm shaking) and 15 µl viral concentrate, and the mixture was poured onto a 20 ml 1.5% bottom agar plate (BD Difco 2216 marine broth). For the enrichment, 100 µl superhost overnight culture was incubated with 20 µl viral concentrates in 10 ml BD Difco 2216 marine broth enrichment cultures at 25°C, and the culture was shaken at 250 rpm for 3 days. These enrichment cultures were passed through a 0.22 µm filter to remove the remaining cells (Minisart High Flow PES, Sartorius), and the filtrate was used with the superhost in an agar overlay for viral plaque isolation as described above. Plaques were picked after one day of incubation at 25°C, and plates were checked for new plaques until two weeks after plating. Viruses were purified three times by restreaking in molten agar and picking individual plaques, and the final stock was prepared by adding 10 ml SM buffer (saline magnesium buffer: 5.8 g*L^-1^ NaCl, 2.0 g*L^-1^ MgSO_4_*7H_2_O, 0.05 M Tris-HCl pH 7.5) to a plate with confluent lysis, incubated for 1 hour and filtered through a 0.22 µm filter (Sartorius). Viral stocks were stored at 4°C, and their taxonomic classification and source of isolation are summarized in Supplementary Table 7.

For sequencing of newly isolated viruses, 10 ml of viral stock was concentrated in 12 ml polypropylene tubes (14 x 89 mm, Beckman Coulter) by ultracentrifugation (Optima XPN-80, Beckman Coulter) in a SW41Ti rotor at 35,000 rpm for 2 h at 4°C. The supernatant was discarded, and the viral pellet redissolved in 100 µl of SM buffer. DNA was extracted with the Pure link Viral RNA/DNA Mini Kit (Invitrogen). Initial lysis was done in ultracentrifugation tubes, and DNA was eluted in PCR-grade water. Viruses were prepared for sequencing with NEBNext® Ultra™ II FS DNA Library Prep Kit for Illumina, pooled equimolarly, and sequenced on an Illumina MiSeq platform (v3 2×300bp) (JMF, University of Vienna). Reads were quality trimmed with BBDuk (v39.01, BBTools, sourceforge.net/projects/bbmap/) using the parameters “ktrim=r k=21 mink=11 hdist=2 tbo=t tpe=t qtrim=rl trimq=15 minlength=50” and interleaved with reformat.sh (BBTools, sourceforge.net/projects/bbmap/). The cleaned reads were assembled with SPAdes (v3.15.5, Bankevivh et al. 2012) with the parameters “-k 31,41,51,61,71,81,91,101,111,121,127 –isolate”. Further polishing and rearrangement were done using Geneious Prime ® (v.2023.2.1. Biomatters Ltd.).

#### Superhost genome sequencing

High molecular weight DNA for long-read sequencing was extracted using the Monarch HMW DNA extraction kit for tissue (NEB), with the specific protocol for Gram-negative bacteria. Extracts were left at 4°C for one week before quantification and fragment estimation. Fragments below 10kb were removed with the Short Read Eliminator kit (PacBio). Samples were diluted and equimolarly barcoded using the SQK-RBK114.96 (Oxford Nanopore Technologies) with the following protocol modifications: we increased the sample input to 80 ng/sample, added 0.2 µl barcode/sample and 0.5 µl of rapid adaptor to the barcoded library. About 18 fmol of a >60 Kb library was loaded on a R10.4.1 flow cell (FLO-PRO114, Oxford Nanopore Technologies) and sequenced for 72h on a Promethion P24 (Oxford Nanopore Technologies) using Minknow (v. 22.10.7, Oxford Nanopore Technologies). We protected the flowcell from light as recommended recently by the manufecturer. Reads were basecalled using Guppy (v. 6.3.9) in super accuracy mode. The Nanopore reads were assembled using flye^58^ (v. 2.9.1) with “–nano-hq,” and polished once with medaka (v. 1.7.2, github.com/nanoporetech/medaka). Reads <1000bp were excluded from the assemblies. Low coverage contigs (<10x) were removed. The medaka model used was “r1041_e82_400bps_sup_g615”. Assemblies were quality controlled using QUAST^59^ (v. 5.2.0) and CheckM^60^ (v. 1.2.2).

Mutant was verified by extracting DNA with the KingFisher using the KingFisher™ Ready DNA Ultra 2.0 Prefilled Plates (ThermoFisher). The library was prepared with the NEBNext Ultra II FS DNA Library Prep Kit for Illumina, barcoded with the NEBNext Unique Dual Index UMI Adaptors DNA Sets 1-4 an sequenced in ½ Illumina NovaSeq S1 flowcell. Reads were quality trimmed with with BBDuk (v39.01, BBTools, sourceforge.net/projects/bbmap/) using the parameters “ktrim=r k=21 mink=11 hdist=2 tbo=t tpe=t qtrim=rl trimq=15 minlength=50”, interleaved with reformat.sh (BBTools, sourceforge.net/projects/bbmap/) and mapped to the reference using the Geneious mapper in highest sensitivity mode.

#### Engineering of a defense-less cloning host (superhost)

Because the *Vibrio* isolates of the Nahant collection naturally contain defense genes, we aimed at creating a strain in which cloned defense and anti-defense genes could be challenged by virus infection assays with greatly reduced interference of chromosomal defense genes. In engineering this strain, we pursued the following strategy.

We previously noticed that identifiable defense genes are associated with large mobile genetic elements (MGEs) in the Nahant vibrios and that, among closely related isolates, genome size scales with the number of mobile genetic elements carrying viral defense genes. Because comparative genomics indicated that *Vibrio cyclitrophicus* 10N.239.312.A01 had one of the smallest genomes in our collection, we chose it for the deletion of all MGEs. We first identified all MGEs in *V. cyclitrophicus* 10N.239.312.A01 by a pangenome-based approach to predict genomic islands using the panrgp workflow of PPanGGOLiN-1.2.105^61^ (*ppanggolin panrgp workflow*). It identified 5 MGEs >5kbp, of which two were identified as prophages using the prediction tools Phaster^62^ (April 15, 2022) and Virsorter2^63^ (v.2.2.4). We then independently identified defense genes using DefenseFinder v.1.0.8^4,64,65^ (March15, 2022), and genomic analysis showed that all defense genes were exclusive to these 5 MGEs. We detected the defense systems Septu, Retron, Gabija, and RM Type IV (Supplementary Data 6). Finally, a reanalysis of defense genes with DefenseFinder v.1.2.2. (March 15, 2024) confirmed that even with the expanded database, no additional defense genes were detectable outside of the knocked-out MGEs.

Because it is still possible that not all defense genes are detected by current database searches, and because defense genes tend to be clustered on MGEs, we aimed to knock out all MGEs containing known defense systems and predicted prophages, in *V cyclitrophicus* 10N.239.312.A01 using the double allelic exchange protocol as described before^3^. All deletions were performed using site-directed mutagenesis, and cloning was executed using the New England Biolabs Gibson Assembly Master Mix following the manufacturer’s protocol. The flanking regions were designed to overlap with the edges of PDE by 60 nt on each side. The length of the flanking regions was calculated as 400 nt + 50 nt multiplied by the length of PDE (in kb) minus 1. Upstream and downstream fragments of the target PDE and the plasmid pSW7848T backbone were PCR-amplified using the specified primers from Supplementary Data 7. The amplicons were treated with DpnI for two hours at 37°C to deactivate the plasmid template before initiating the Gibson assembly reaction. All PCR products were verified with gel electrophoresis, purified using PCR Cleanup kit (Qiagen), and quantification of DNA concentration using a Qubit HS kit (Invitrogen) according to manufacturer instructions. Subsequently, three fragments were mixed in 0,05 pmol concentrations (the volumes for the 20 mcl Gibson assembly reaction were calculated online NEBioCalculator) and incubated at 50°C for 60 minutes. 5-10 mcl of Gibson assembly mixture were used for transformation into chemically competent *E.coli* pi3813 cells prepared according to the protocol described in^66^. After one hour incubation at 37°C, the cells were plated in 10^0^, 10^-1^, and 10^-2^ dilutions on selective Luria Broth agar (LBA) plates containing 0.3mM dT and 25mcg/ml chloramphenicol, and were incubated at 37°Covernight. The colonies were screened for insert using Check1_ins_pSW_for and Check1_ins_pSW_rev primers from Supplementary Data 7 using Phusion polymerase (NEB, Phusion High Fidelity DNA Pol, M0530L) polymerase for inserts >5kb (PCR thermocycling conditions were as follows: initial denaturation at 98 °C for 30 sec; 30 cycles of 98 °C for 10 sec, 60 °C for 20 sec, 72 °C for 15sec/kb depending on the length of the fragment; final annealing at 72 °C for 5 min; hold at 10 °C), and OneTaq (New England Biolabs) polymerase for inserts <5kb (PCR thermocycling conditions were as follows: initial denaturation at 94 °C for 30 sec; 30 cycles of 94 °C for 20 sec, 60 °C for 30 sec, 68 °C for 1mic/kb depending on the length of the fragment; final annealing at 68 °C for 5 min; hold at 10 °C). The inserts of a right size were purified using QIAquick PCR Purification Kit (Qiagen) and Sanger sequenced. Lastly, the plasmid DNA was purified, confirmed via Sanger sequencing, and transformed into chemically competent *E. coli* β3914 described in^66^ to serve as a plasmid donor for conjugation.

Conjugation was carried out by mating assays as described previously^3^ with some modifications. First, the donor: recipient ratio was adjusted to 3:1. The mating was performed directly from the overnight cultures of the donor and recipient. Mating spots were resuspended in 50 mcl of SM buffer and 100 mcl of 0, −1, −2 and −3 dilutions were spread onto TSB1 plates supplemented with Cm and incubated at 25°C for two days.

The resulting *V. cyclitrophicus* 10N.239.312.A01 strain, which we named “superhost” lacked any known defense systems in its genomes, making it suitable for cloning external defense systems without interference from its inherent defense mechanisms as well as isolation of viruses unbiased by resident defense systems.

#### Cloning of defense systems

For all cloning and expression, we used the shuttle vector pMRB-PlacGFP^67^, which contains an oriVR6Kg origin of replication, a chloramphenicol resistance cassette, and a green fluorescent protein (GFP) gene under a constitutive lac promoter. For cloning of defense systems, we amplified each defense operon from selected bacterial isolates of the Nahant collection for which defense and anti-defense interactions were predicted. To include the native promoter, we designed the PCR primers to comprise a region 500 nt upstream of the start codon (Supplementary Data 7), used Phusion polymerase (NEB, Phusion High Fidelity DNA Pol, M0530L) for amplification, and cloned the product into pMRB-PlacGFP replacing the GFP gene using the NEB Gibson assembly mixture kit.

#### Cloning of candidate anti-defense genes

The anti-defense candidates were cloned on the same plasmid as the respective defense system, downstream of the defense system and under a constitutive lac promoter. For this, candidate anti defense genes were first commercially synthetized (until not specified below) and cloned in the shuttle vector pBBR1MCS5 congaing oriTpBBR1 origin of replication and gentamicin resistance cassette instead of the *lacZ* gene (so that they would appear under an inducible promoter after cloning) by ThermoFisher. All open reading frames encoding candidate anti defense proteins were cloned along with 100 bp of upstream sequence, and their sequences are provided in Supplementary Table 9. The primers pBBR1_BamHI_F and pBBR1_MluI_R (Supplementary Data 7) annealing to the 25-long flanking regions of the cloned anti-defense candidates+promoter with restriction sites for BamHI and MluI were designed to amplify the insert from flanking regions. Each DNA fragment was previously checked for lack of BamHI and MluI restriction sites. The DNA of each anti-defense candidate was amplified from the pBBR1_antidefense_candidate source using Phusion polymerase and was hydrolyzed with FD Thermo Fisher BamHI and MluI enzymes according to the manufacturer’s instructions. The hydrolyzation products were PCR purified and were used for further cloning. The plasmids containing defense systems were hydrolyzed with FD Thermo Fisher BamHI and MluI and used for ligation reaction with the respective anti defense-containing fragment with T4 ligase according to the standard protocol. 5 µl of each ligation reaction was used to transform chemically competent *E.coli* pi3813 cells as described above, and the colonies were screened for the insert of a right size with the primers pMRB-PlacGFP_ins_1_F and pMRB-PlacGFP_ins_1_R using Phusion polymerase (NEB, Phusion High Fidelity DNA Pol, M0530L) polymerase for the insert longer than 5kb (PCR thermocycling conditions were as follows: initial denaturation at 98 °C for 30 sec; 30 cycles of 98 °C for 10 sec, 60 °C for 20 sec, 72 °C for 15sec/kb depending on the length of the fragment; final annealing at 72 °C for 5 min; hold at 10 °C), and OneTaq polymerase (NEB, OneTaq Quick-Load DNA polymerase, M0509L) for the inserts shorter than 5kb (PCR thermocycling conditions were as follows: initial denaturation at 94 °C for 30 sec; 30 cycles of 94 °C for 20 sec, 60 °C for 30 sec, 68 °C for 1mic/kb depending on the length of the fragment; final annealing at 68 °C for 5 min; hold at 10 °C). All constructions were verified using Sanger sequencing. The resulting plasmids were transformed into chemically competent *E. coli* beta3914 cells as described above, which were used as a donor to transfer the plasmids into *Vibrio* recipient cells via conjugation.

For all cloned defense and anti-defense pairs, we evaluated whether the overexpression of anti-defense proteins impacted the expression of the defense system by conducting quantitative MS/MS counts for both defense and anti-defense proteins. In every case, the overexpression of anti-defense proteins did not alter the expression of the defense system.

#### Plaque assays/Phage challenge assays

To test the efficiency of defense or anti-defense, we determined the efficiency of plating using 10-fold viral dilutions. Bacterial strains were prepared for agar overlay plating by streaking out from glycerol stocks onto 2% streak agar plates (BD Difco 2216 marine broth), and allowed to grow for one days at room temperature. Strains were then inoculated into 5 ml of BD Difco 2216 marine broth in a 15 ml culture tubes and incubated 24 h at room temperature shaking at 250 rpm. Viral lysates were propagated on their host strains by plating stock lysates onto agar overlays as described before^3^. Plaque assays were performed as follows. A 250 µl volume of overnight bacterial culture was combined with 10 ml of molted 0,4% top agar (BD Difco 2216 marine broth) preheated to 42°C, the mixture was then transferred to the surface of 10-cm square 1,5% bottom agar plates (BD Difco 2216 marine broth) and incubated for 1 hour at room temperature. 10-fold serial dilutions of the phages were prepared in SM buffer for each tested phage, and 2-µl drops of each dilution were applied onto the bacterial layer. Plates were incubated at room temperature overnight. Plaque forming units (PFUs) were determined by counting clearings in the bacterial lawn after 24h incubation, and lysate titer was calculated as PFUs per ml. In cases where individual plaques were not discernible, a faint lysis zone covering the drop area was considered equivalent to 10 plaques as described^28^. Efficiency of plating was assessed by comparing plaque assay results between control bacteria (cloning host with GFP-containing plasmid) and those harboring the cloned defense system or defense system and candidate anti-defense gene.

#### Proteomics Workflow: Sample preparation, LC-MS measurements, and Data analysis

To determine whether the cloned defense genes were expressed in a similar fashion whether they were expressed alone or in combination with anti-defense genes, we carried out LC-MS analysis.

5 ml of *Vibrio* bacteria were grown overnight at 25°C with 250 rpm shaking in BD Difco 2216 marine broth medium. Overnight cultures were freshly diluted in 5 ml of BD Difco 2216 marine broth medium (1:100 volume) and the cultures were grown to 0,5 OD600. 5 ml of the culture was pelleted and immediately frozen in liquid nitrogen. For sample preparation, the bacterial cell pellets were subjected to acetone precipitation at a ratio of 1:5 (v/v). After precipitation, the air-dried pellets were solubilized in lysis buffer (2M Urea, 100mM ammonium bicarbonate, protease inhibitor). Following sonication for 40 seconds on ice, each sample lysate was centrifuged at 16,000g and 4°C for 10 minutes. The supernatants were transferred to clean tubes, and then protein concentrations were determined using the Bio-Rad Bradford protein assay. A minimum of 30 µg of protein was applied for alkylation with 500 mM Iodoacetamide (room temperature, shaking at 700 rpm, 60 min in the dark), and for disulfide reduction with 500 mM of DTT (room temperature, shaking at 700 rpm, 15 min). In-solution on-bead digestion of proteins was performed using Immobilized trypsin (Promega, sample-enzyme solution ratio of 94 (v/v)) with an overnight incubation of 16 hours at 37°C.

Peptide desalting was carried out using OMIX C18 solid-phase extraction (SPE) tips from Agilent Technologies. After equilibration of the stage-tip with 100% methanol, the tip was washed twice with formic acid (FA) solution (0.1% FA in high purity water (MilliQ)) (v/v). Peptide elution was performed with 100% methanol. The peptide concentration was determined using Pierce™ Quantitative Colorimetric Peptide Assay (Thermo Scientific). The peptides were dried out using a SpeedVac and were dissolved in a 0.1% FA solution (v/v) [FA in high purity water (MilliQ)].

1µg of peptides was separated by an online reversed-phase (RP) HPLC (Thermo Scientific Dionex Ultimate 3000 RSLC nano LC system) connected to a benchtop Quadrupole Orbitrap mass spectrometer (Q-Exactive Plus, Thermo Scientific). The online separation was performed on an Easy-Spray analytical column (PepMap RSLC C18, 2 μm, 100 Å, 75 μm i.d. × 50 cm, Thermo Scientific) with an integrated emitter. The Column was heated at 55°C. The flow rate was set to 300 nL/min.

For LCMS analysis the LC gradient employed a two-and-a-half-hour gradient. It started at 2% and gradually increased to 35% buffer B (v/v) [79.9% ACN, 0.1% FA, 20% Ultra high purity (MilliQ)] over 130 minutes, followed by a transition to 80% buffer B over 5 minutes. The buffer A (v/v) was 0.1% FA in high purity water (MilliQ). LC eluent was introduced into the mass spectrometer through an Easy-Spray ion source (Thermo Scientific). The emitter was operated at 1.9 kV.

Two different MS methods were employed: one using a DDA for controls, and the other utilizing a combined DDA-Parallel reaction monitoring (PRM) method for analysing the plasmid peptide samples. All methods were measured in positive ion mode. For DDA method, a top 10 data-dependent acquisition (DDA) was employed. The full mass spectrum was acquired at a resolution of 70,000 at m/z 200 [AGC target at 3e6, maximum injection time (IT) of 120 ms and a scan range 200-1200 (m/z)]. Following the MS scan, the MS/MS scan was performed at 17,500 resolution at m/z 200 [Automatic Gain Control (AGC) target at 1e5, 1.6 m/z isolation window and maximum IT of 100 ms]. For MS/MS fragmentation, normalized collision energy (NCE) for higher energy collisional dissociation (HCD) was set to 28%. Dynamic exclusion was set at 50 s. Unassigned and +1, +6, +7, +8 and > +8 charged precursors were excluded. The intensity threshold was set to 5.0e3. The isotopes were excluded.

The combined Full MS with PRM, the fullMS was set to 70,000 resolution at m/z 200 [AGC target at 1e6, maximum injection time (IT) of 100 ms and a scan range 200-1200 (m/z)]. The PRM (tandem MS) method was set to 35,000 resolution at m/z 200 [AGC target at 1e6, 2.0 m/z isolation window, maximum injection time (IT) of 100 ms]. For MS/MS fragmentation, stepped normalized collision energy (NCE) for higher energy collisional dissociation (HCD) was set to 25%, 28% and 30%. The fragment ion mass window was set to ±5 ppm. The corresponding precursor ion masses (m/z) of 10-14 selected doubly and triply changed peptides were set up in the inclusion list for each plasmid peptide sample.

For the combined Full MS with PRM method, the full MS was set to 70,000 resolution at m/z 200 [AGC target at 1e6, maximum injection time (IT) of 100 ms and a scan range 200-1200 (m/z)]. The PRM method was set to a resolution of 35,000 at m/z 200 [AGC target at 1e6, 2.0 m/z isolation window, maximum injection time (IT) of 100 ms]. For MS/MS fragmentation, stepped normalized collision energy (NCE) for higher energy collisional dissociation (HCD) was applied, set at 25%, 28%, and 30%. The fragment ion mass window was set to ±5 ppm. Additionally, an inclusion list was created for each plasmid peptide sample, containing corresponding precursor ion masses (m/z) of an average 40 selected doubly and triply charged peptides.

All of the raw data for *Vibrio cyclitrophicus* were searched against the proteome sequences obtained from the UniProtKB *Vibrio cyclitrophicus* complete proteome database (downloaded on March 8th, 2024), as well as custom-made plasmid protein database, using MaxQuant software (version 2.3.1.0). Similarly, all of the raw data for *Vibrio lentus* were searched against the proteome sequences obtained from the UniProtKB *Vibrio lentus* complete proteome database (downloaded on March 8th, 2024), along with a custom-made plasmid protein database, using MaxQuant software (version 2.3.1.0).

The tandem mass spectra were analyzed by the Andromeda search engine. A precursor ion tolerance of 20 ppm and a fragment ion tolerance window of 0.5 Da were selected. There was a maximum of two missed cleavages, with static carbamidomethylation of cysteine residues (+57.021 Da), and variable oxidation of methionine residues (+15.995 Da). The false discovery rate (FDR) was set to 0.01 (1%) at both protein and peptide levels. Matches between the runs were selected for identification, and contaminants and reversed data were filtered out from protein group lists. The identified proteins were selected based on at least one peptide (unique and razor) using the custom-made plasmid database. For identifications based on large-databases, at least one unique peptide was required.

## Data availability

New bacterial and phage genomes deposited with this work are included under the Nahant Collection of NCBI BioProject with accession number PRJNA328102. All sequence accession numbers are provided in Supplementary Data 4.

## Code availability

The code for all analysis detailed in the main text and in the following sections, including genome identifiers and scripts, will be available upon publication.

## Acknowledgement

We would like to acknowledge the Life Science Compute Cluster at the University of Vienna for providing access to high-performance computing clusters. We thank the Joint Microbiome Facility of the Medical University of Vienna and the University of Vienna for their support with sequencing. We would like to acknowledge Professor Zbyszek Otwinowski for providing access to high-performance computing clusters and Marcin Pietrzycki for assistance with massive computations (on the architectural side of Deep Learning algorithms and its scaling to large datasets). We thank Javier Dubert Perez for providing plasmids and for advice and guidance with double-allelic exchange experiments. We thank Jacob Bobonis for discussions and comments at various stages of the study. This work was supported by the Austrian Science Fund (FWF) [doi.org/10.55776/COE7], the Simons Foundation (Life Sciences Project Award-572792) (MFP), the Austrian Science Fund (FWF) grant (Grant-DOI 10.55776/P33891 from Prof. Dr. Verena Ibl (Cell Biology in Crop Seeds (CELBICS) group)) (MS), a Marie Curie postdoctoral fellowship (Project 101063227 PHAGECOUNTER) (AL).

## Author’s contributions

A.L. and M.F.P. developed the concept and designed the study. M.F. and developed the model and performed predictions with input from A.L. S.P. conducted the comparative genome analysis. A.L., M.W and N.B performed knock-outs to construct superhost. A.L. and M.W performed cloning and plaque assays. S.M. performed experimental verification of interaction of *Vibrio lentus* and its phages. K.S. performed prophage search and provided R script for heatmaps. L.A.S designed the proteomic mass spectrometry workflow, supervised sample preparation, performed LCMS analysis. processed and interpreted the data, and wrote the Methods section for Proteomics mass spectrometry. M.S. performed sample preparation for LC-MS analysis. A.L, S.P., N.B. designed figures, with inputs from M.F.P. A.L., S.P. and M.F.P. prepared the manuscript with input from all coauthors.

## Supplementary materials

### Extended Data Figures

**Extended Data Figure 1.**
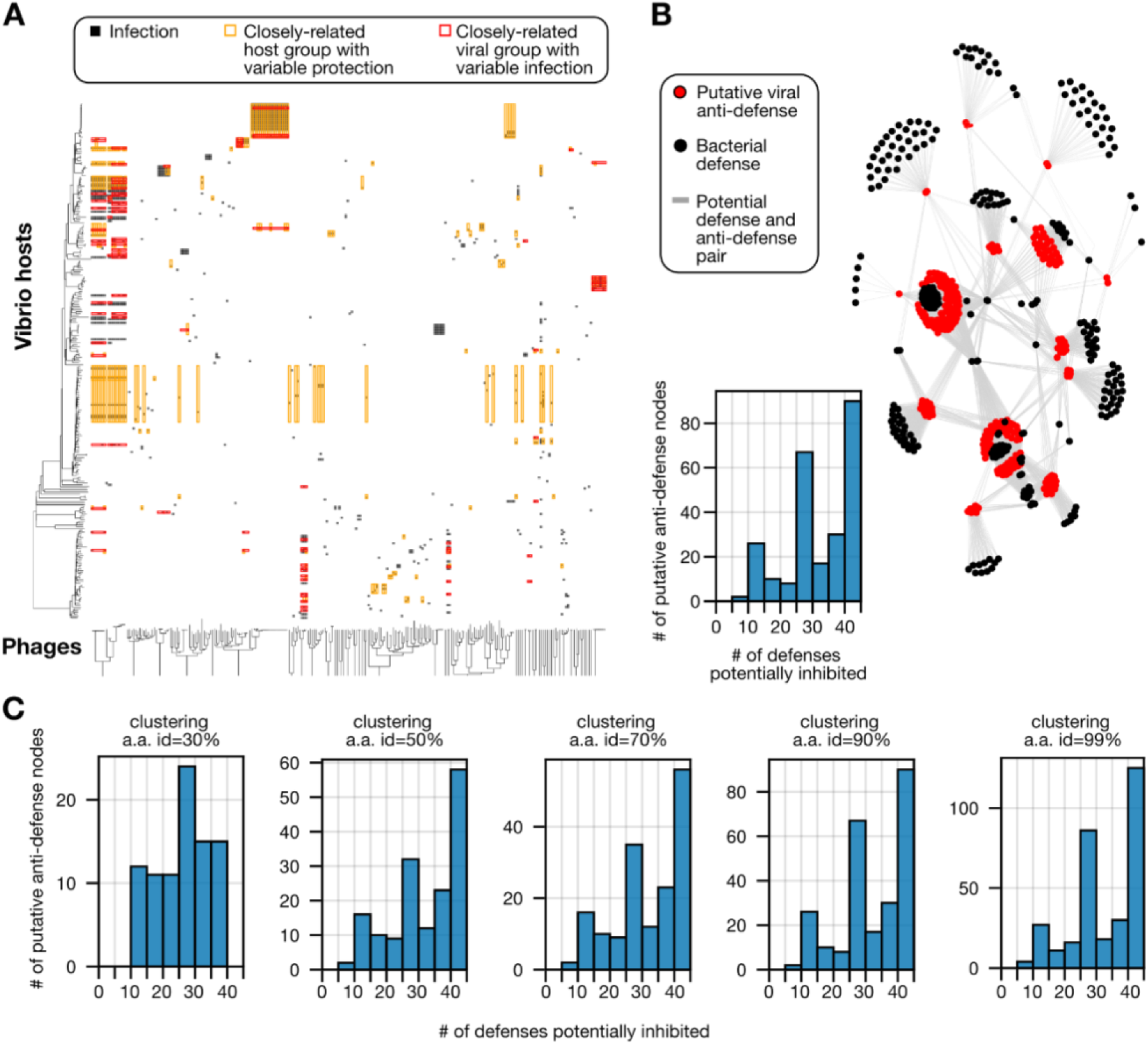
Associating bacterial defenses to putative viral anti-defense genes is limited by the high number of defenses co-occurring in bacterial genomes. **a,** The Nahant cross-infection matrix with relevant closely related viral (phage, x-axis) and bacterial groups (y-axis) highlighted. The trees represent FastME dendrograms built from ANI distances. Entries in the matrix are marked with a black box if a particular phage can infect and lyse a particular bacterial host. Groups of closely related phages (average ANI ≥ 0.95) that differentially infect a particular bacterial host are marked on the matrix with a red rectangle surrounding the relevant infection data of the phage group and the host. Closely related bacteria (Average ANI ≥ 0.95) that are differentially protected from a single phage are marked in the same way using golden rectangles. Putative anti-defenses are identified within each phage group, potential inhibitory interactions are identified across all genomes (methods). **b,** Network showing connections between bacterial defenses and the putative viral anti-defenses that inhibit them based on a comparative genomic analysis of the Nahant cross-infection matrix. Putative anti-defense genes (red nodes) in viral genomes were identified as gene families (clustered at 90% identity) found only in viruses infecting a single bacterial host, compared to their close relatives that do not infect that host (methods). Bacterial defenses (black nodes) in each host were identified using DefenseFinder, and links are drawn between putative anti-defenses and the defenses encoded in the infected host. The histogram shows the degree distribution of each anti-defense, i.e., how many defenses it putatively inhibits. Identifying specific inhibition of defenses by anti-defenses would require extensive trial and error genetics approaches. **c,** putative anti-defense node degree histograms for different protein clustering amino-acid (a.a) similarity thresholds (indicated at the top of each histogram). The distributions are very skewed towards high values for all thresholds, and there are no clear-cut cases of defense and anti-defense pairs in any threshold.

**Extended Data Figure 2.**
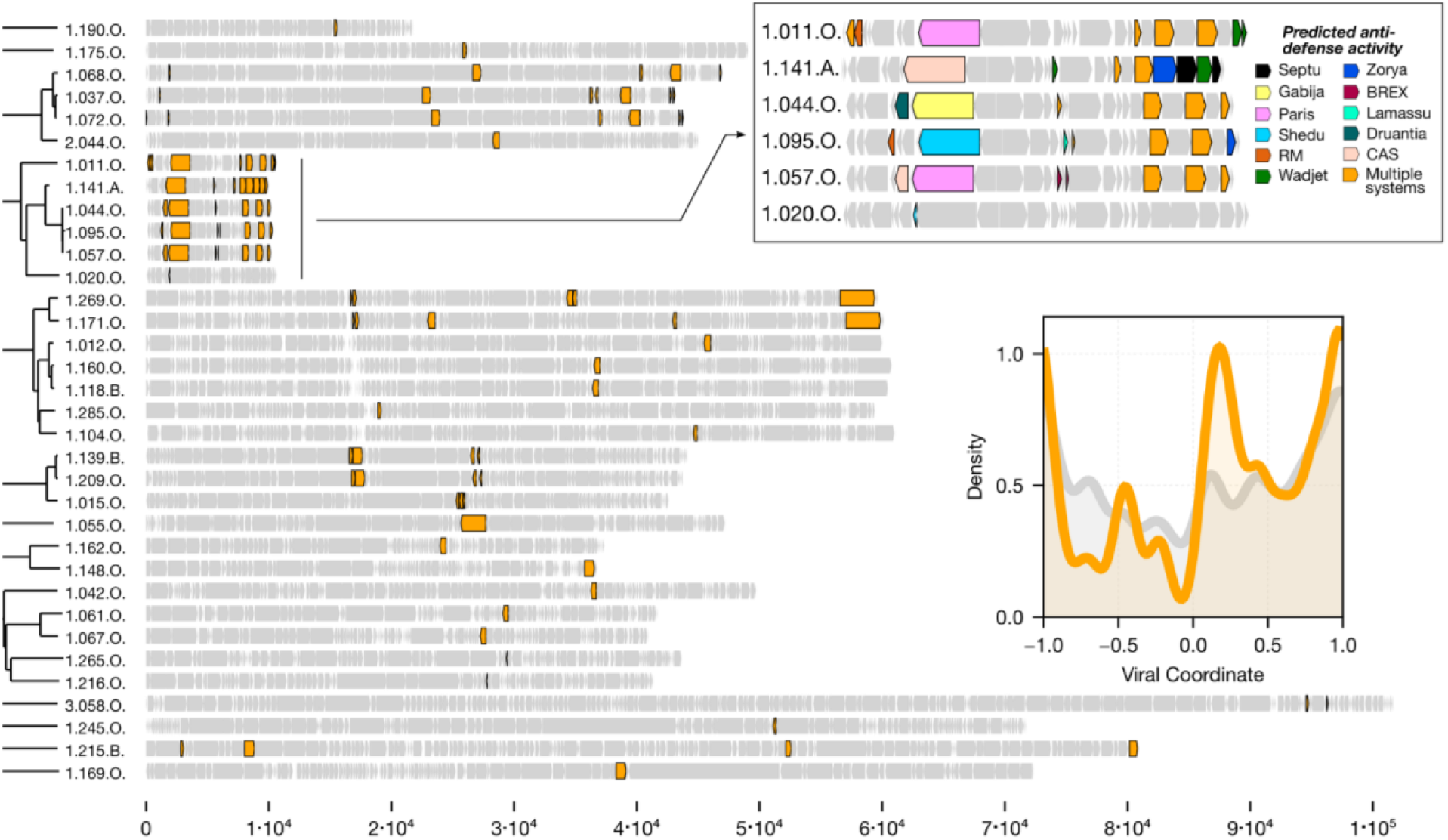
Mapping of putative anti defenses identified by VirMAD onto viral genomes of the Nahant cross-infection matrix. The tree on the left was built using fastME^68^ based on ANI values between full viral genomes. Clades that are disconnected have no detectable homology to viruses from other clades. Only viruses that contained homologues (>90% a.a similarity) of putative anti-defenses are shown. Each row represents a viral genome, starting at the viral terminase small subunit with arrows representing genes to scale as marked in the bottom. The highlighted viruses have short genomes, which contain many putative anti-defenses with different specificities. The density plot on the right shows the normalized position of putative anti-defenses (orange curve) in the viral genomes compared to all genes (gray curve).

**Extended Data Figure 3.**
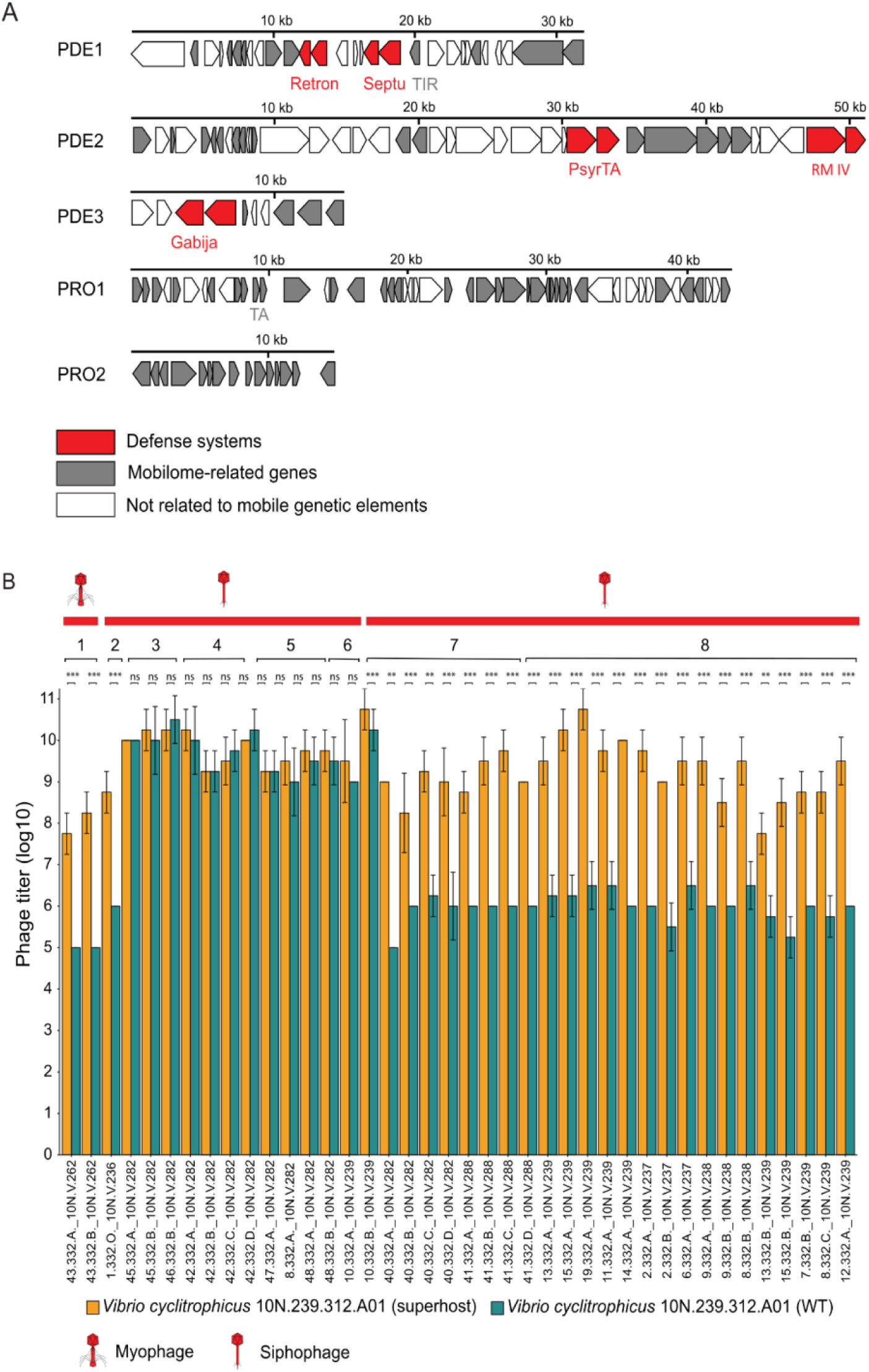
Deletion of phage defense systems in the superhost results in increased phage sensitivity. **a,** Creation of the superhost strain by deletion of mobile genetic elements from *Vibrio cyclitrophicus* 10N.239.312.A01, chosen because they either contained predicted defenses or represented prophages. Arrow gene diagrams drawn to scale (annotations are in Supplementary Table 10). **b,** Comparison of titers of 40 viruses on wild type (teal) and superhost (orange) showing increased susceptibility of the superhost. Numbers under red bars represent clusters of viruses based on genome similarity (Supplementary Table 16). Data represent average of three independent replicates with standard deviations (s.d.) shown as vertical error bars.

**Extended Data Figure 4.**
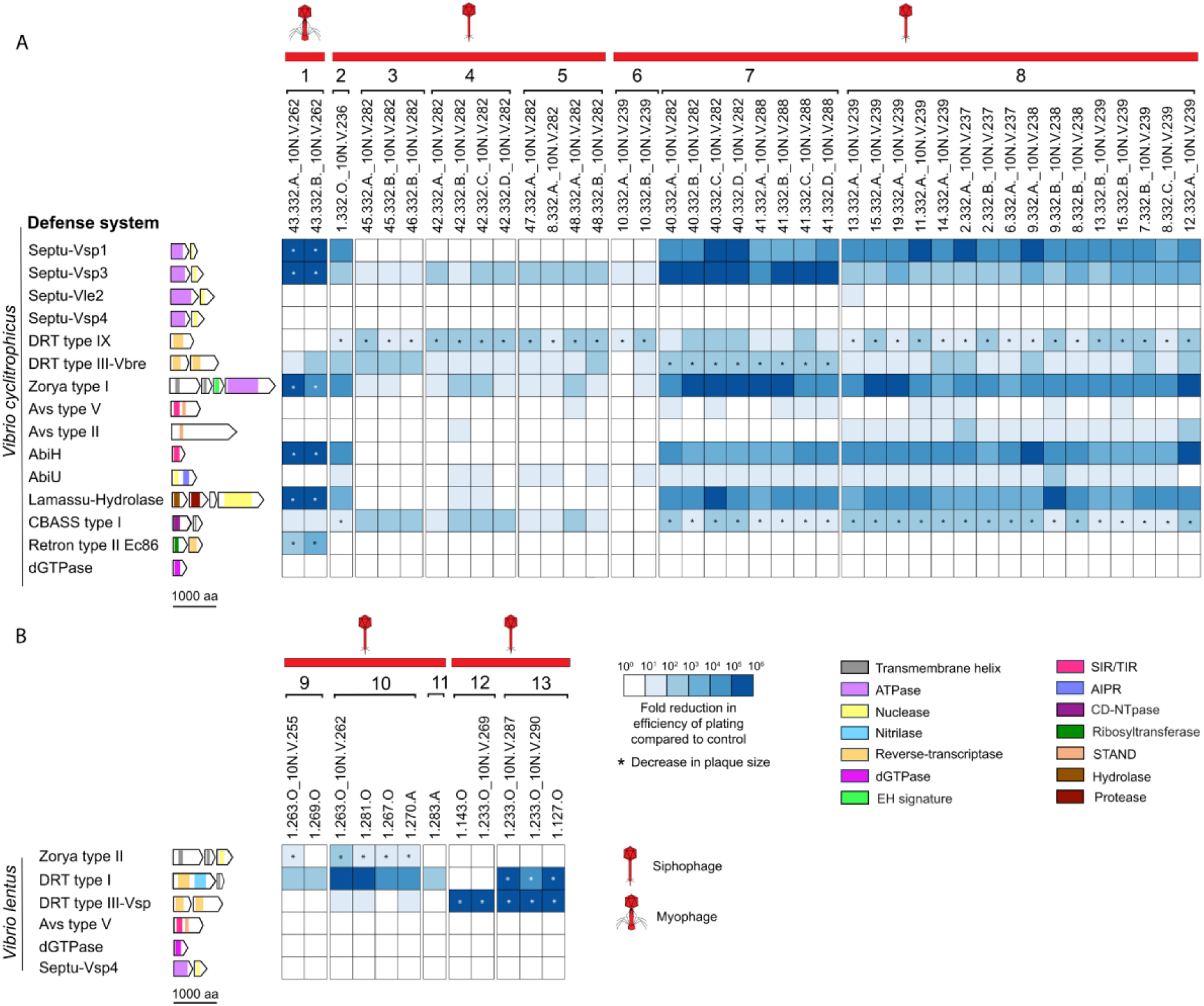
Fold-protection conferred by all experimentally verified defense systems, Defenses expressed in **a,** the *Vibrio cyclitrophicus* superhost and **b,** *Vibrio lentus* Δ3 (methods). The strength of protection is shown as a heatmap where values correspond to the fold change in efficiency of plating (measured in serial dilution plaque assays) compared to an unprotected host (GFP expressing plasmid) for each virus. Numbers above virus identifiers represent clusters of viruses based on genome similarity (Supplementary Table 16). Data represent the average of three replicates (examples of raw data shown in Supplementary Fig. 1). Asterisks (*) indicate significant reduction in plaque size. Arrow gene diagrams with predicted protein domains in color are drawn to scale.

**Extended Data Figure 5.**
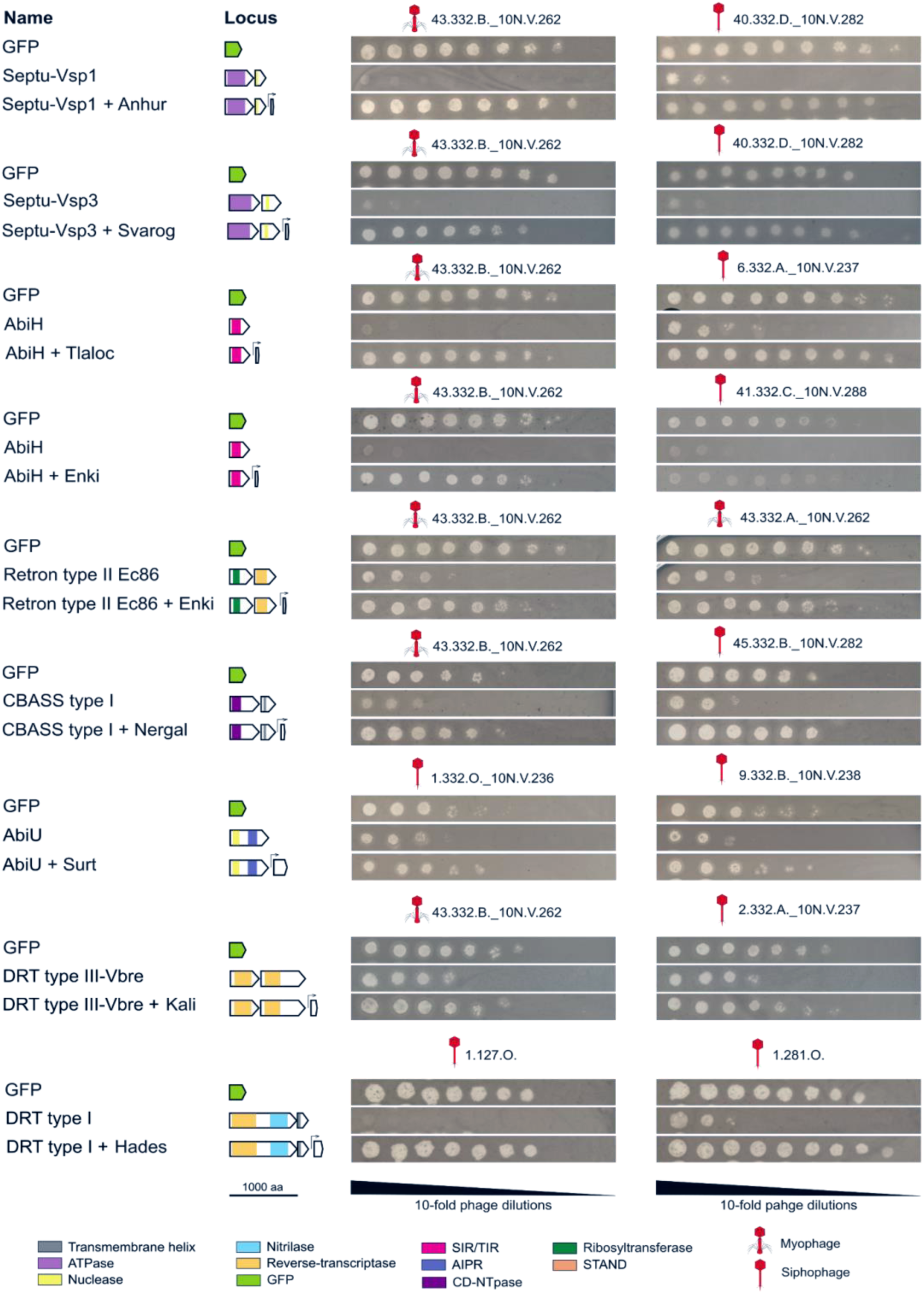
Drop spots of 10-fold serially diluted viruses on hosts expressing GFP (control), defense systems or paired defense systems and anti-defense proteins. This figure accompanies Fig. 2 and Extended Fig. 4. For each paired defense and anti-defense, the two viruses are shown for which the defense system provided the strongest protection. Plaque assays are representative of three independent replicates. Arrow gene diagrams with predicted protein domains in color are drawn to scale.

**Extended Data Fig. 6.**
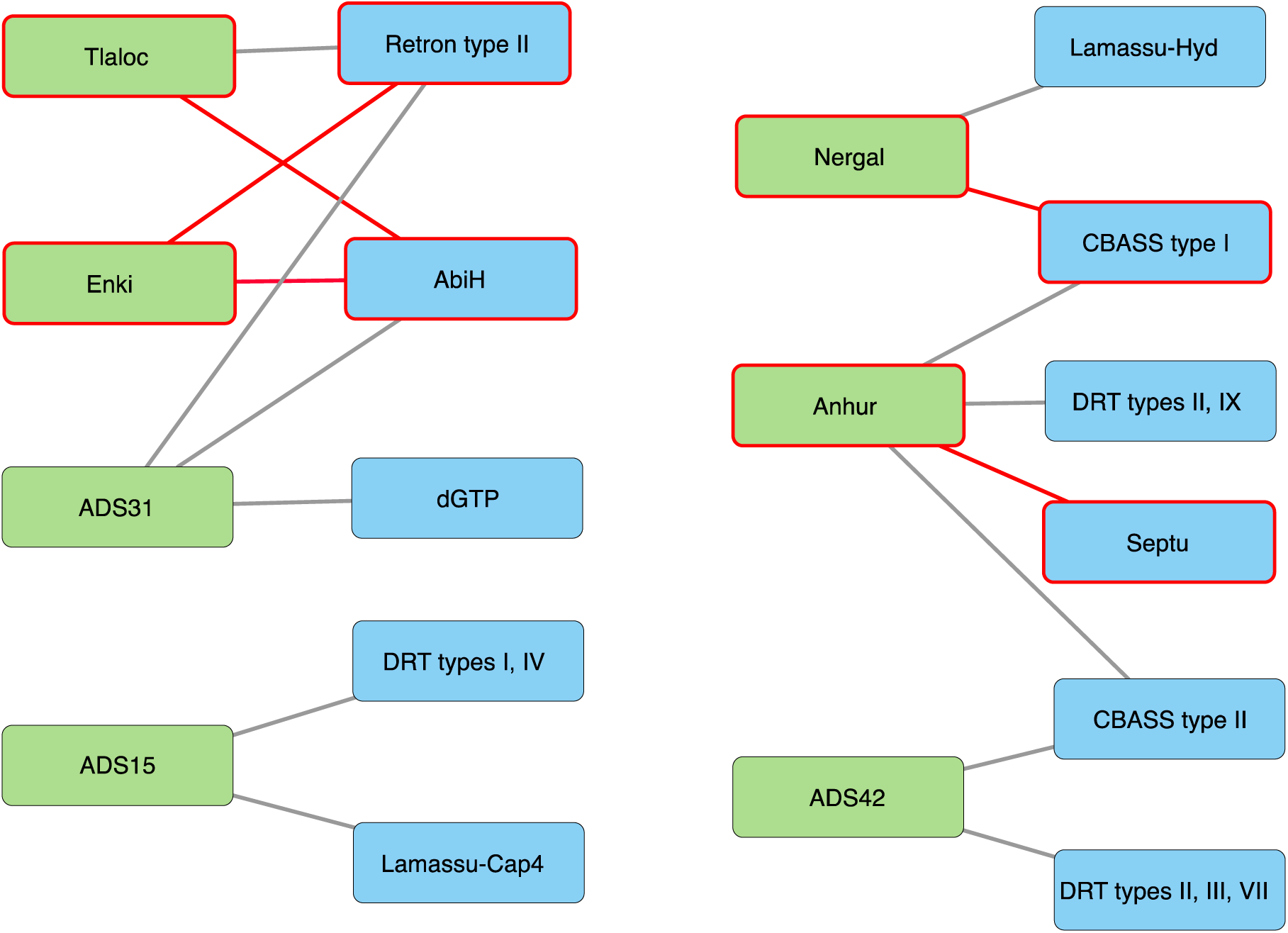
Anti-defense proteins with multiple non-homologous targets, and defense systems with multiple non-homologous inhibitors. The network shows highly scored (>80) VirMAD predictions of defense-anti-defense pairs for ten defense systems chosen for experimental verifications. Only anti-defense proteins with two or more predicted types of defense systems are shown in the network. Putative anti-defense proteins (green boxes) are clustered at 95% sequence similarity, while defense systems (blue boxes) were classified into subtypes using DefenseFinder (see Supplementary Tables 1 and 4). Red lines correspond to defense-anti-defense pairs which were experientially verified.

### Supplementary Figures

**Supplementary Fig. 1.**
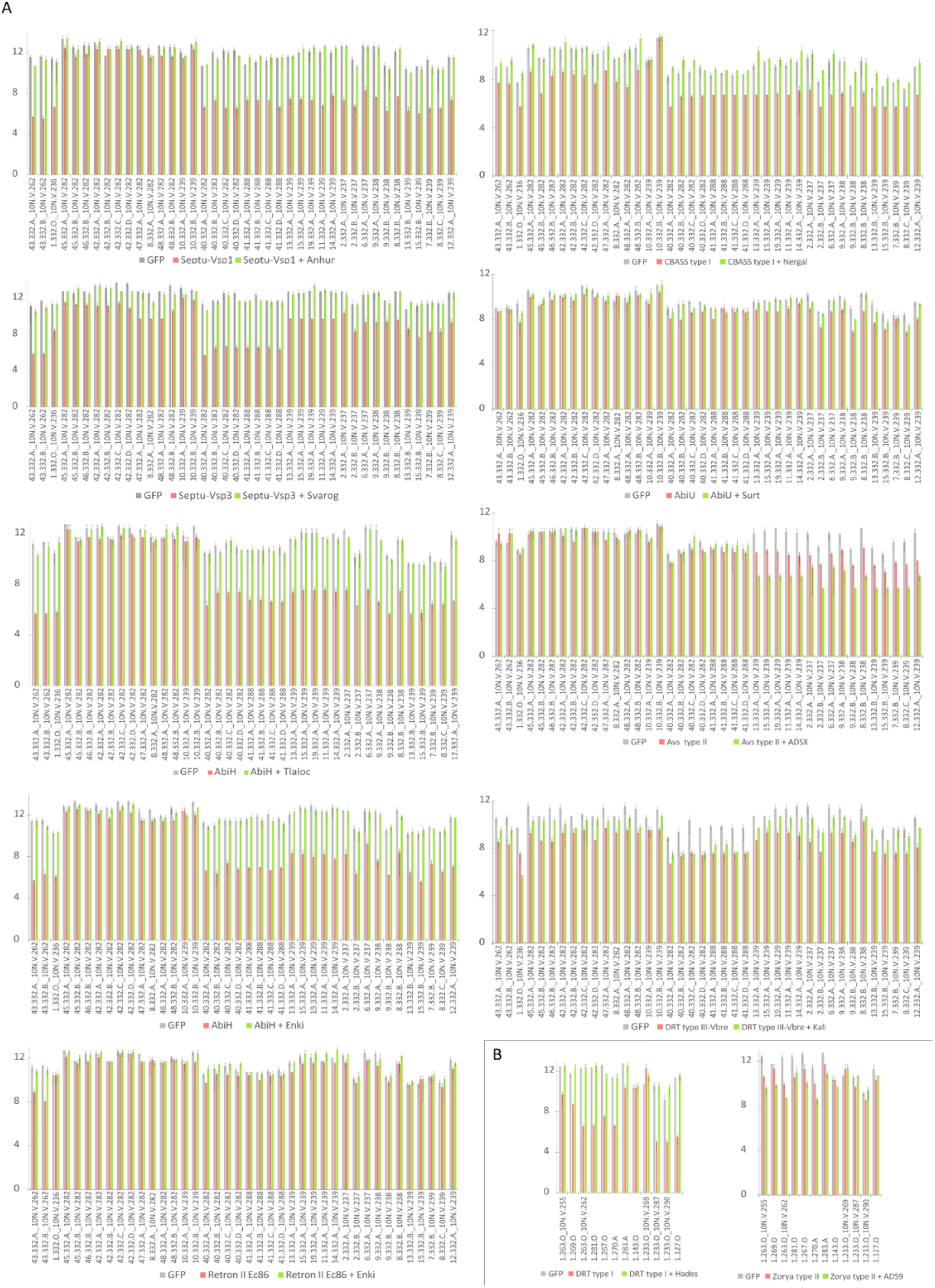
Raw data for fold-protection calculation. This figure accompanies Fig. 2 and Extended Fig. 4. Defense systems expressed in **a,** *Vibrio cyclitrophicus* superhost and **b,** *V. lentus* Δ3 (see Methods). Fold-protection data for all verified pairs of defense systems and anti-defense proteins were obtained for 40 viruses for the superhost strain and 12 viruses for the Δ3 strain. Plaque-forming units were measured for each virus on hosts containing either a GFP-expressing control plasmid (grey), a defense-expressing plasmid (red), or a plasmid expressing both the defense system and anti-defense protein (green). Data represent an average of three independent replicates with standard deviations (s.d.) shown as a vertical bars. All defense systems were expressed from their native promoters, while anti-defense genes were under the control of an inducible *PlacI* promoter. Virus titers were measured after 24 hours of incubation at 25°C.

**Supplementary Fig. 2.**
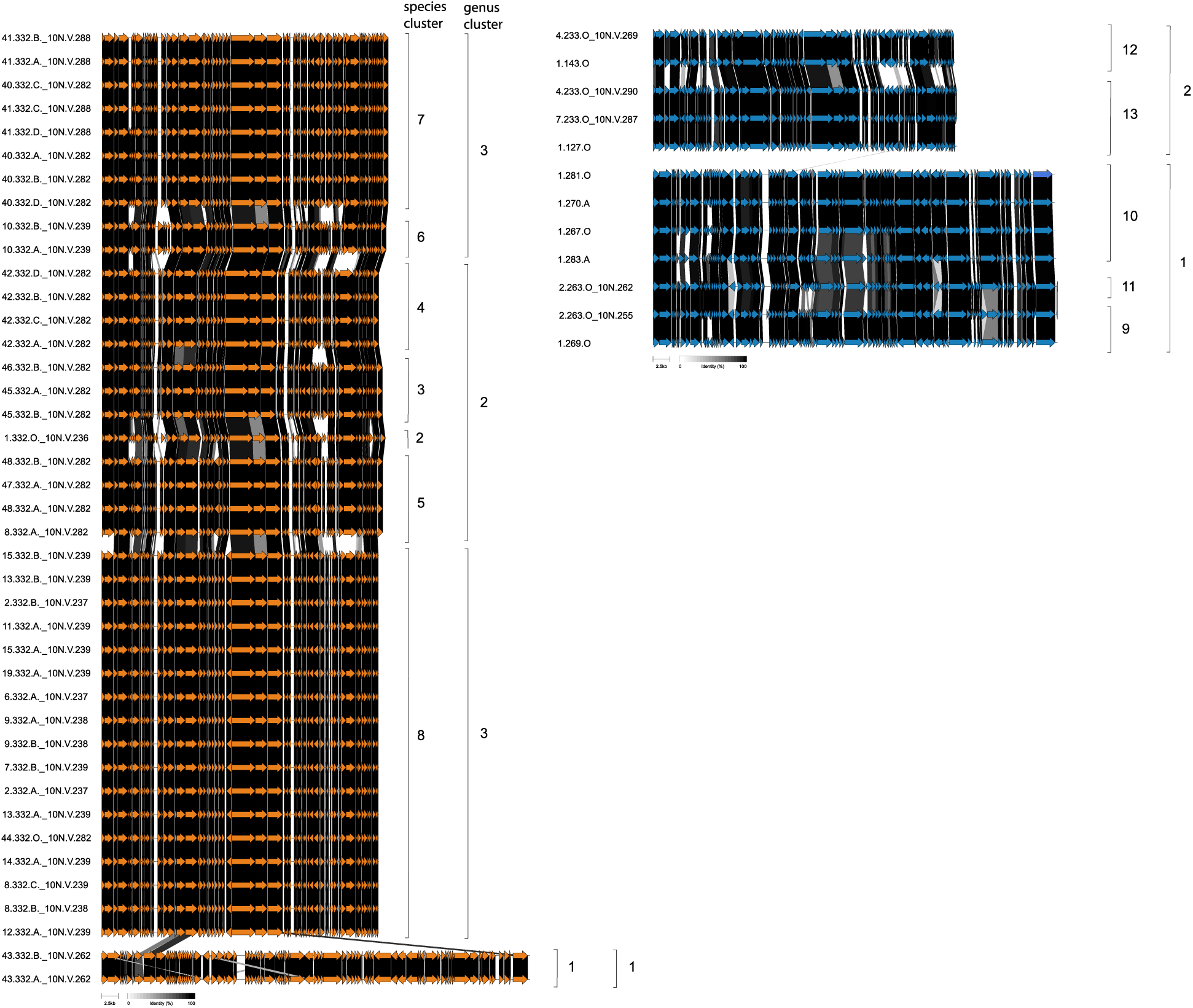
Genomic diversity of *Vibrio cyclitrophicus* and *V. lentus* viruses used in plaque assays. Genomic comparison of the 40 viruses infecting *Vibrio cyclitrophicus* superhost (orange) and the 12 viruses infecting *Vibrio lentus* Δ3 (blue) used in experiments underlying the data in Fig. 2 and Extended Fig. 4. Grey shading indicates the percent identity between homologous proteins. Species and genus cluster predictions (Viridic) are shown on the right.

**Supplementary Fig. 3.**
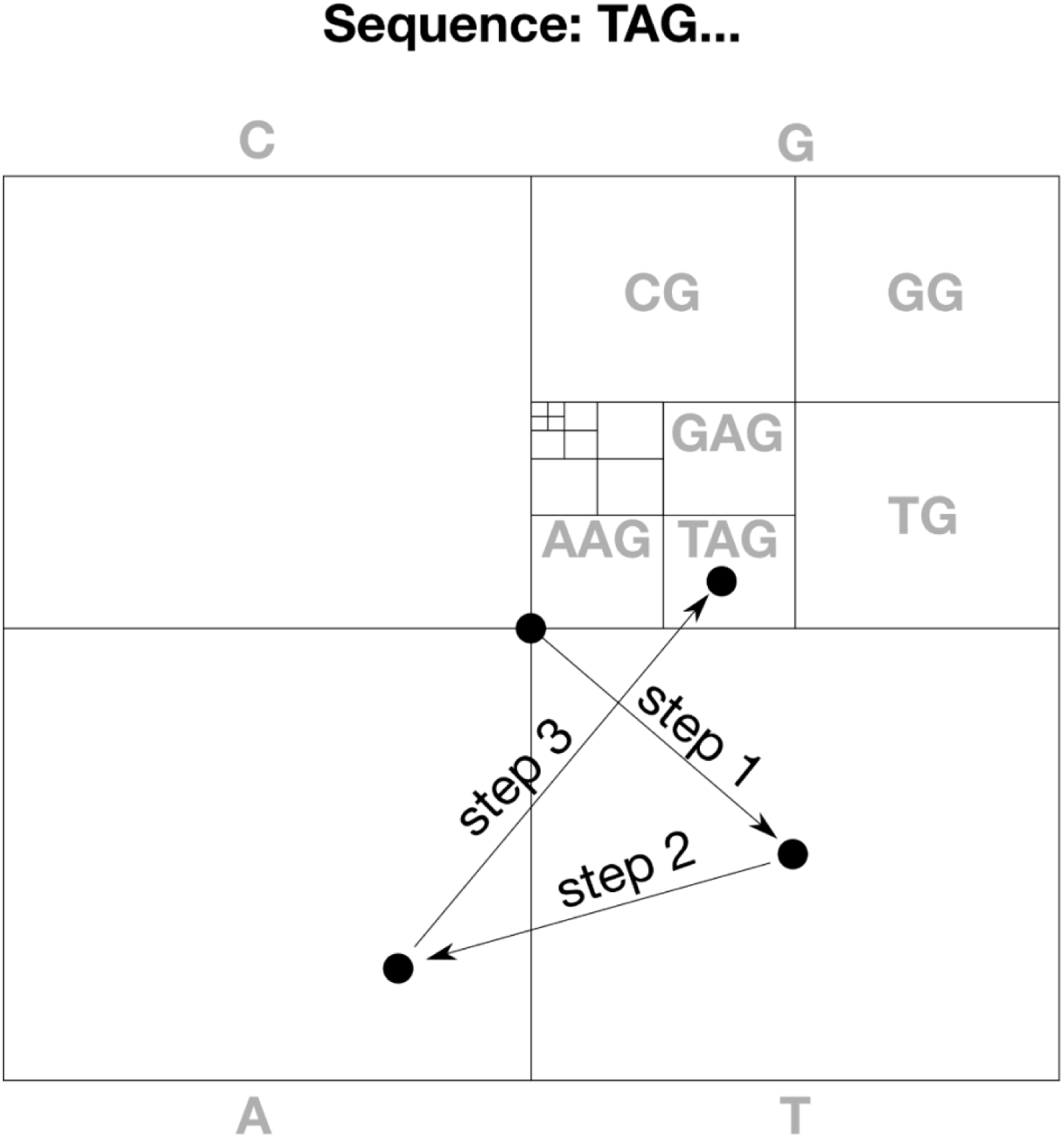
Chaos Game Representation of nucleotide sequences. To visually represent nucleotide sequences, each corner of a square is assigned to a nucleotide. Starting out in the middle of the square, the sequence is processed one base at a time, each time going from the current location halfway to the corner representing the nucleotide being processed. This creates fractal regions corresponding to increasingly long k-mers, as represented in the figure with smaller and smaller squares that divide the total area of the figure. An illustration is provided for the first 3 steps in a sequence that starts with the nucleotides TAG.

**Supplementary Fig. 4.**
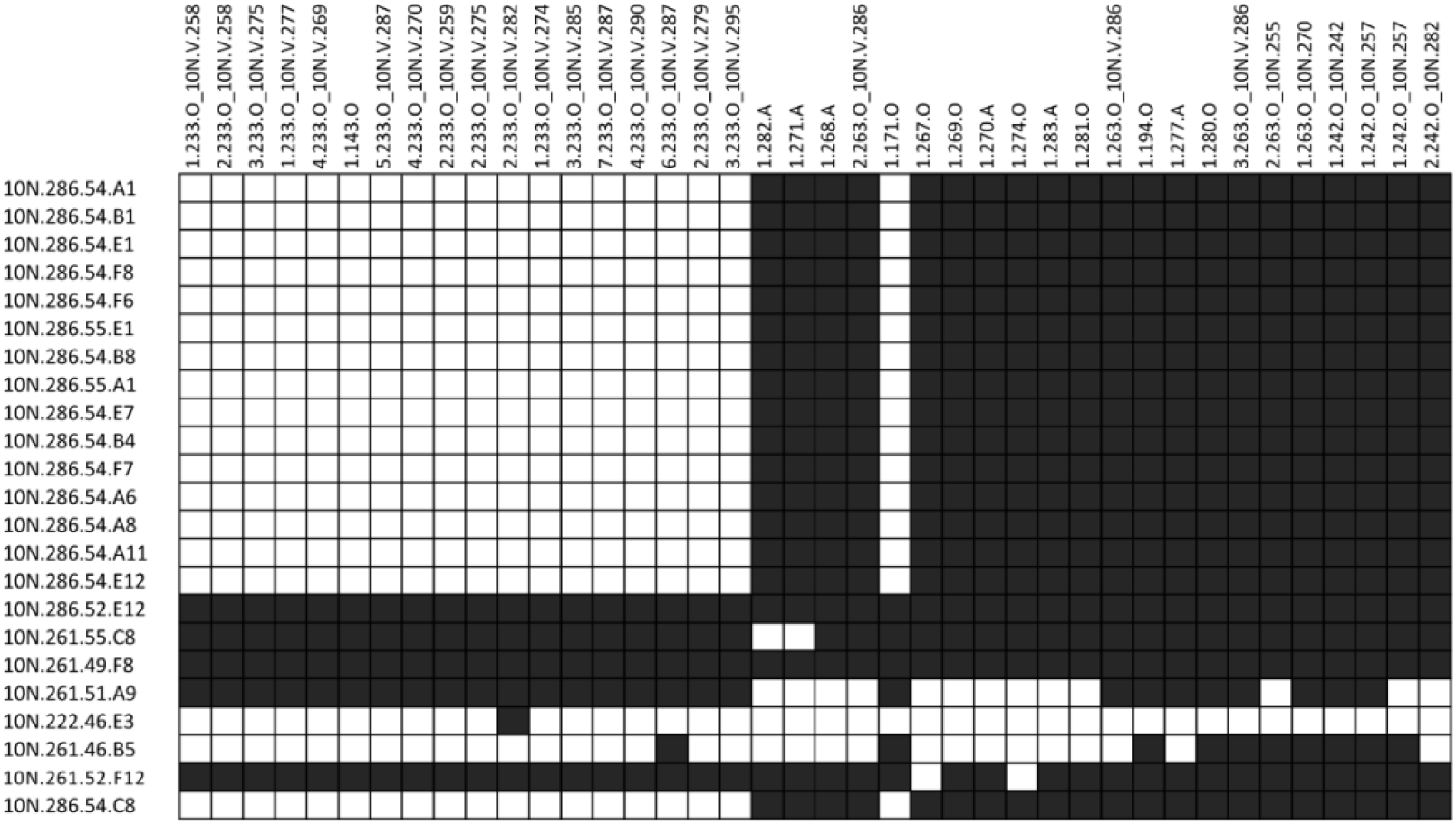
Virus-host cross-infection matrix supplementing the Nahant matrix. Forty viruses were isolated on 5 *Vibrio lentus* strains (methods) and tested against an extended set of 25 *V. lentus* isolates. Positive interactions (lysis) are shown as dark-grey square and negative interactions as white squares. Bacterial strains are ordered based on phylogeny of concatenated single copy ribosomal protein genes; viruses are ordered based on manual sorting of VIRIDIC genus-level trees. This additional matrix was used in the VirMAD predictions.

### Supplementary Tables

**Supplementary Table 1.**
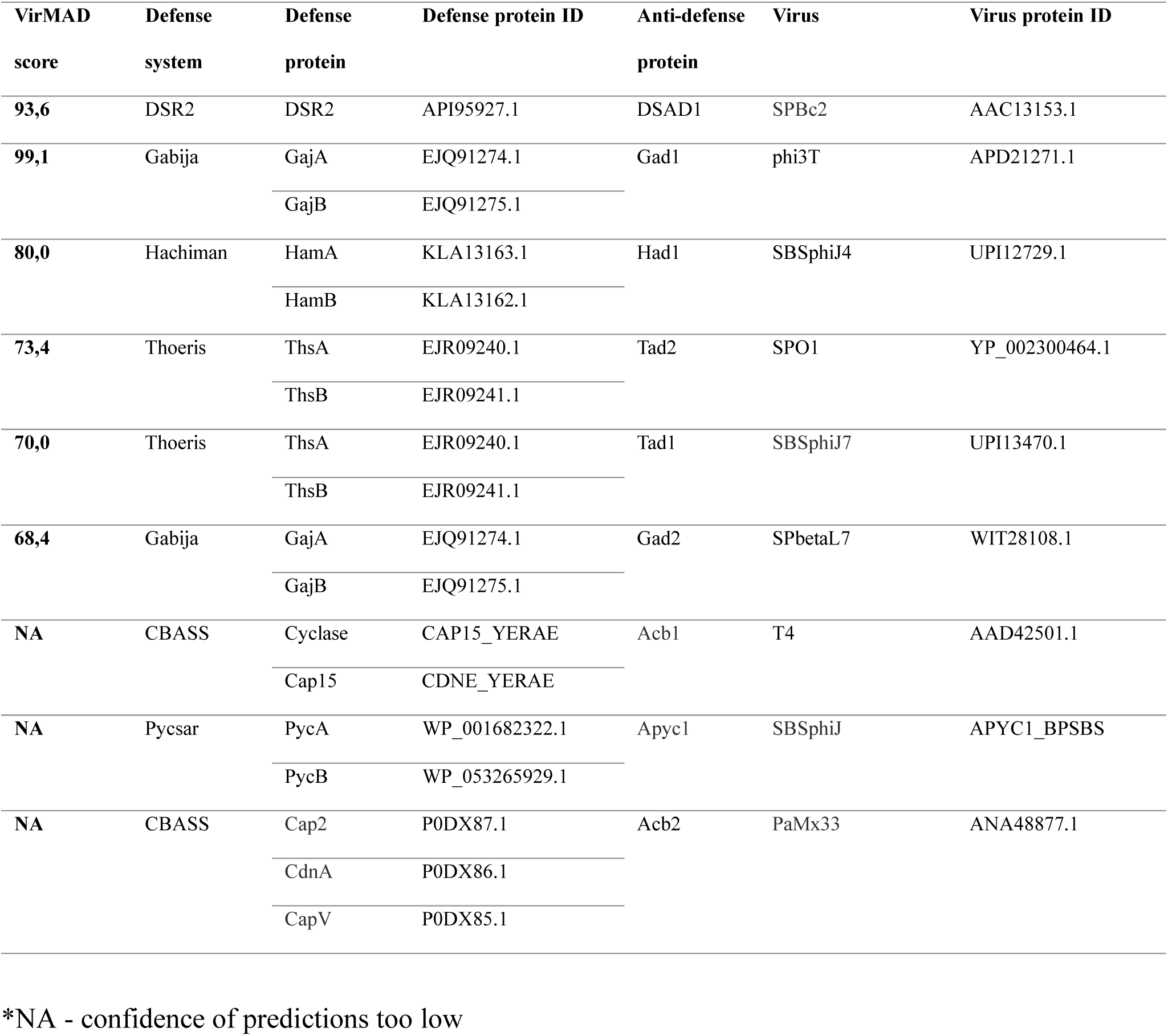
Nine known anti-defense proteins used as positive controls for model predictions.

**Supplementary Table 2.** Relative abundance of defense systems in 758 *Vibrio* genomes from the Nahant collection.

| Defense system type | % relative abundance | VirMAD prediction | Experimental verification of defense phenotype |
| --- | --- | --- | --- |
| **Restriction-modification** | 35,9 | YES |  |
| **dGTPase** | 13,6 | YES | YES |
| **Septu** | 7,1 | YES | YES |
| **CRISPR** | 4,9 | YES |  |
| **Gabija** | 4,3 | YES |  |
| **CBASS** | 3,8 | YES | YES |
| **Retron** | 3,7 | YES | YES |
| **Wadjet** | 2,6 | YES |  |
| **DRT** | 2,4 | YES | YES |
| **Dnd** | 2,0 | YES |  |
| **AbiH** | 1,4 | YES | YES |
| **Rst_PARIS** | 1,3 | YES |  |
| **Zorya** | 1,3 | YES | YES |
| **Hachiman** | 1,1 |  |  |
| **SspBCDE** | 1,1 |  |  |
| **Kiwa** | 1,0 |  |  |
| **Shedu** | 0,9 | YES |  |
| **Lit** | 0,8 |  |  |
| **Druantia** | 0,8 | YES |  |
| **Viperin** | 0,8 |  |  |
| **DarTG** | 0,7 |  |  |
| **dCTPdeaminase** | 0,7 |  |  |
| **Avs** | 0,7 | YES | YES |
| **Gao_Ppl** | 0,6 |  |  |
| **BREX** | 0,6 | YES |  |
| **AbiU** | 0,5 | YES | YES |
| **Thoeris** | 0,5 |  |  |
| **Dsr** | 0,5 |  |  |
| **Gao_Hhe** | 0,4 |  |  |
| **Rst_3HP** | 0,4 |  |  |
| **Lamassu** | 0,4 | YES | YES |
| **Gao_Qat** | 0,4 |  |  |
| **PrrC** | 0,3 |  |  |
| **AbiEii** | 0,3 |  |  |
| **Gao_Her** | 0,3 |  |  |
| **Gao_Ape** | 0,3 |  |  |
| **Gao_Tmn** | 0,3 |  |  |
| **RADAR** | 0,3 |  |  |
| **Rst_DprA-PPRT** | 0,2 |  |  |
| **Rst_HelicaseDUF2290** | 0,2 |  |  |
| **Rst_DUF4238** | 0,2 |  |  |
| **BstA** | 0,1 |  |  |
| **DISARM** | 0,1 |  |  |
| **Gao_Iet** | 0,1 |  |  |
| **Gao_RL** | 0,1 |  |  |
| **Nhi** | 0,1 |  |  |
| **Rst_TIR-NLR** | 0,1 |  |  |
| **Gao_Mza** | 0,1 |  |  |
| **Gao_Upx** | 0,1 |  |  |
| **Rst_Hydrolase-3Tm** | 0,1 |  |  |

**Supplementary Table 3.**
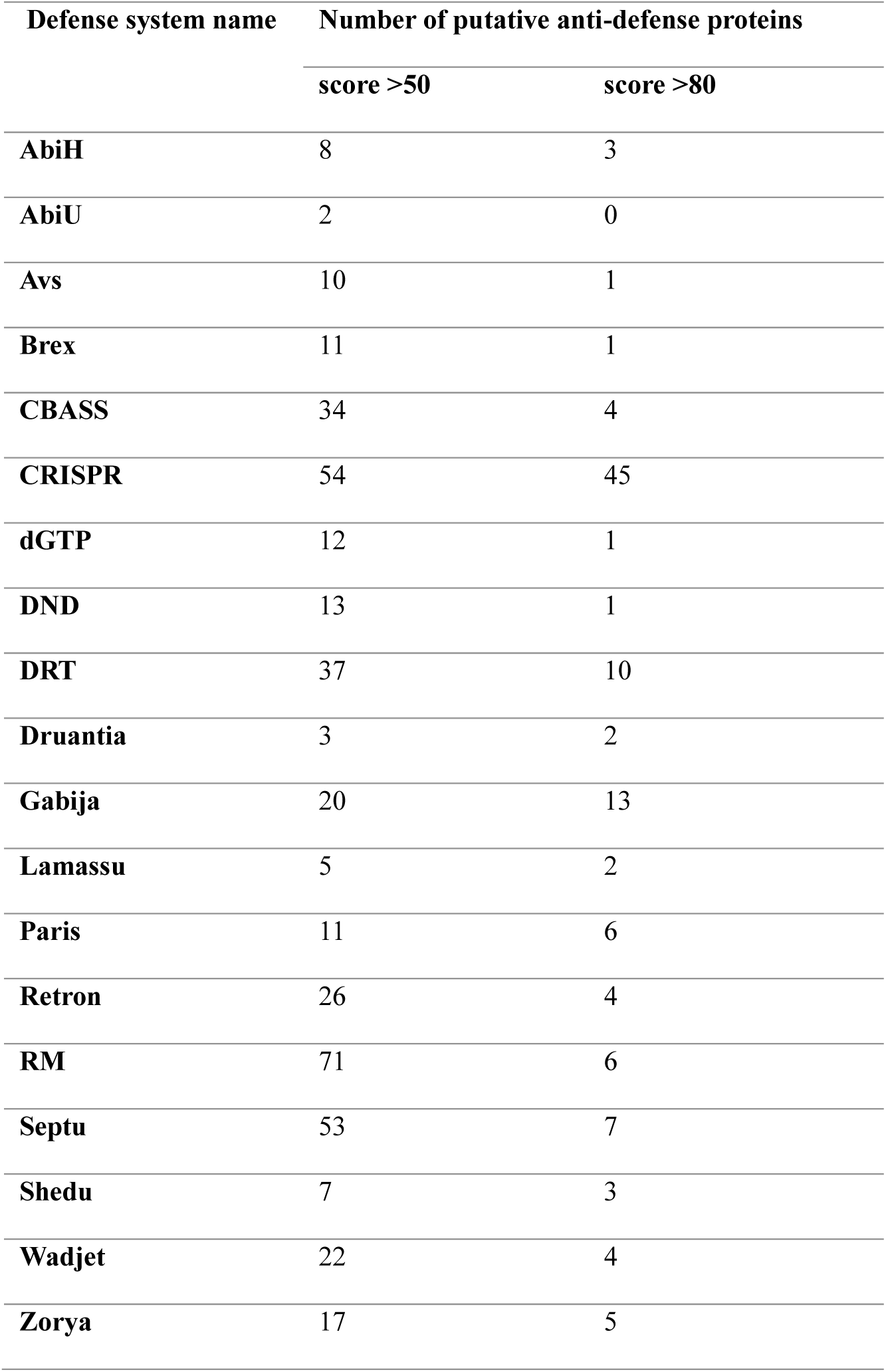
Number of putative anti-defense proteins predicted by the VirMAD model for 19 defense system types.

| Defense system name | Number of putative anti-defense proteins |  |
| --- | --- | --- |
|  | score >50 | score >80 |
| **AbiH** | 8 | 3 |
| **AbiU** | 2 | 0 |
| **Avs** | 10 | 1 |
| **Brex** | 11 | 1 |
| **CBASS** | 34 | 4 |
| **CRISPR** | 54 | 45 |
| **dGTP** | 12 | 1 |
| **DND** | 13 | 1 |
| **DRT** | 37 | 10 |
| **Druantia** | 3 | 2 |
| **Gabija** | 20 | 13 |
| **Lamassu** | 5 | 2 |
| **Paris** | 11 | 6 |
| **Retron** | 26 | 4 |
| **RM** | 71 | 6 |
| **Septu** | 53 | 7 |
| **Shedu** | 7 | 3 |
| **Wadjet** | 22 | 4 |
| **Zorya** | 17 | 5 |

**Supplementary Table 4.**
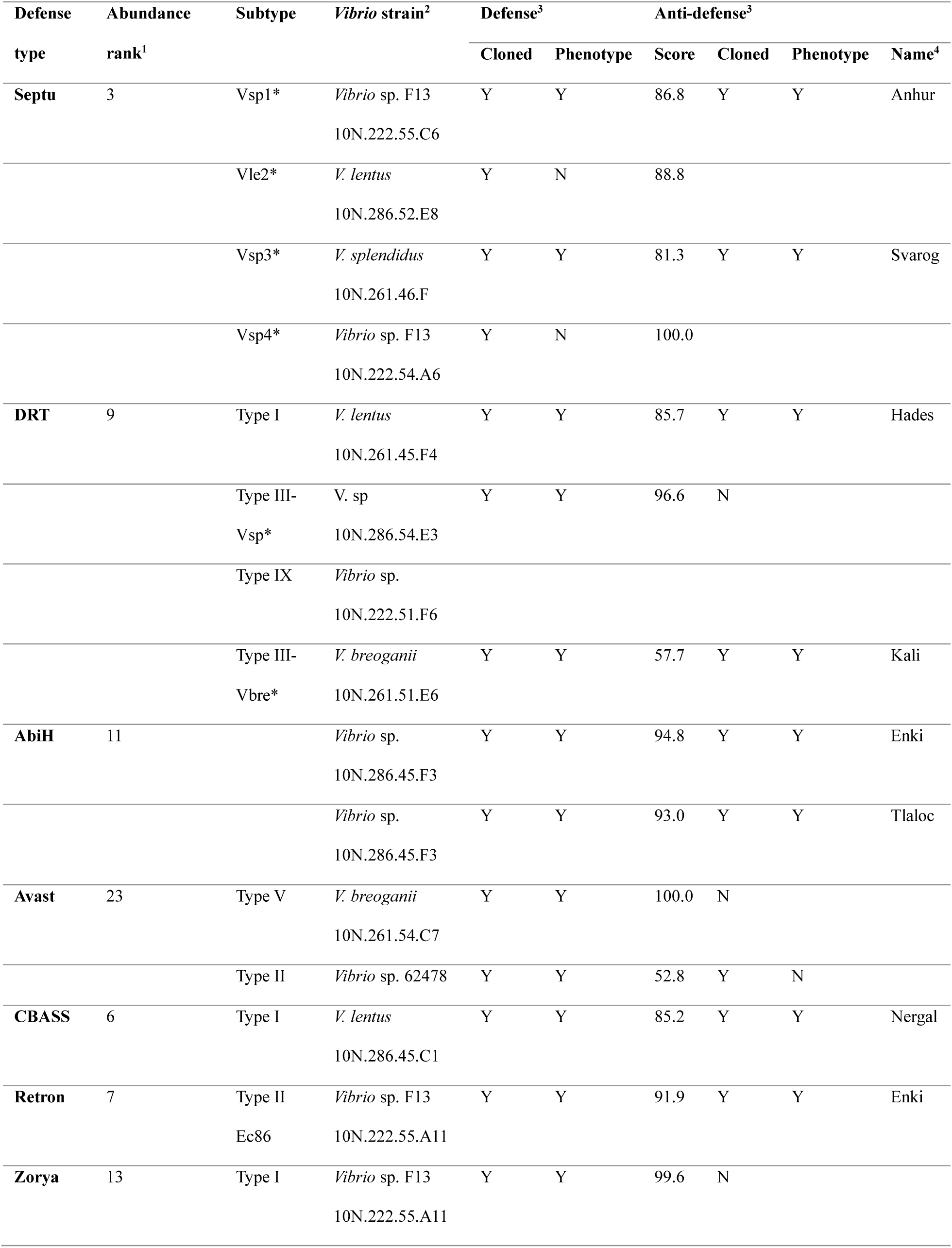
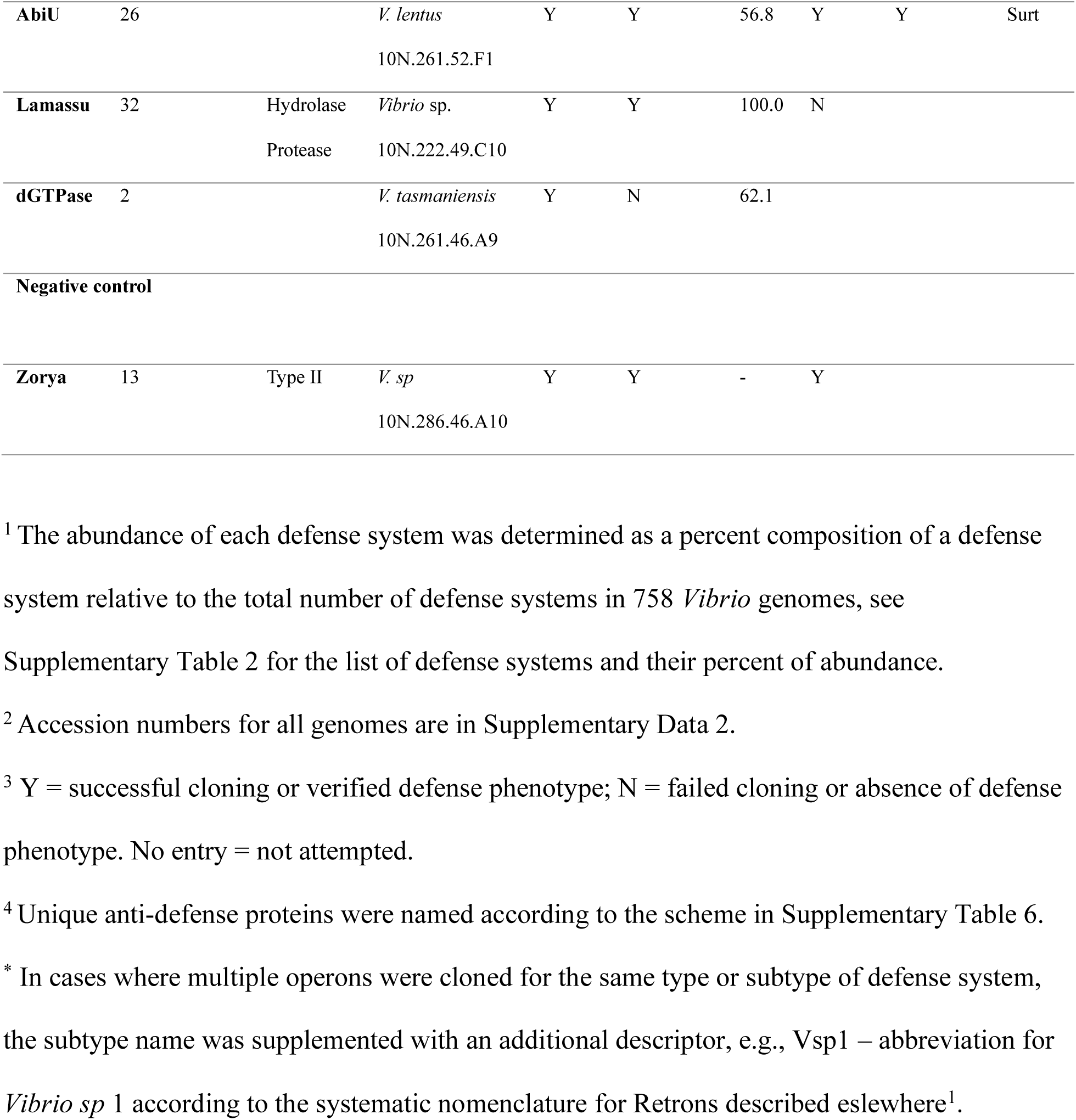
Cloned defense systems and putative anti-defense proteins.

**Supplementary Table 5.**
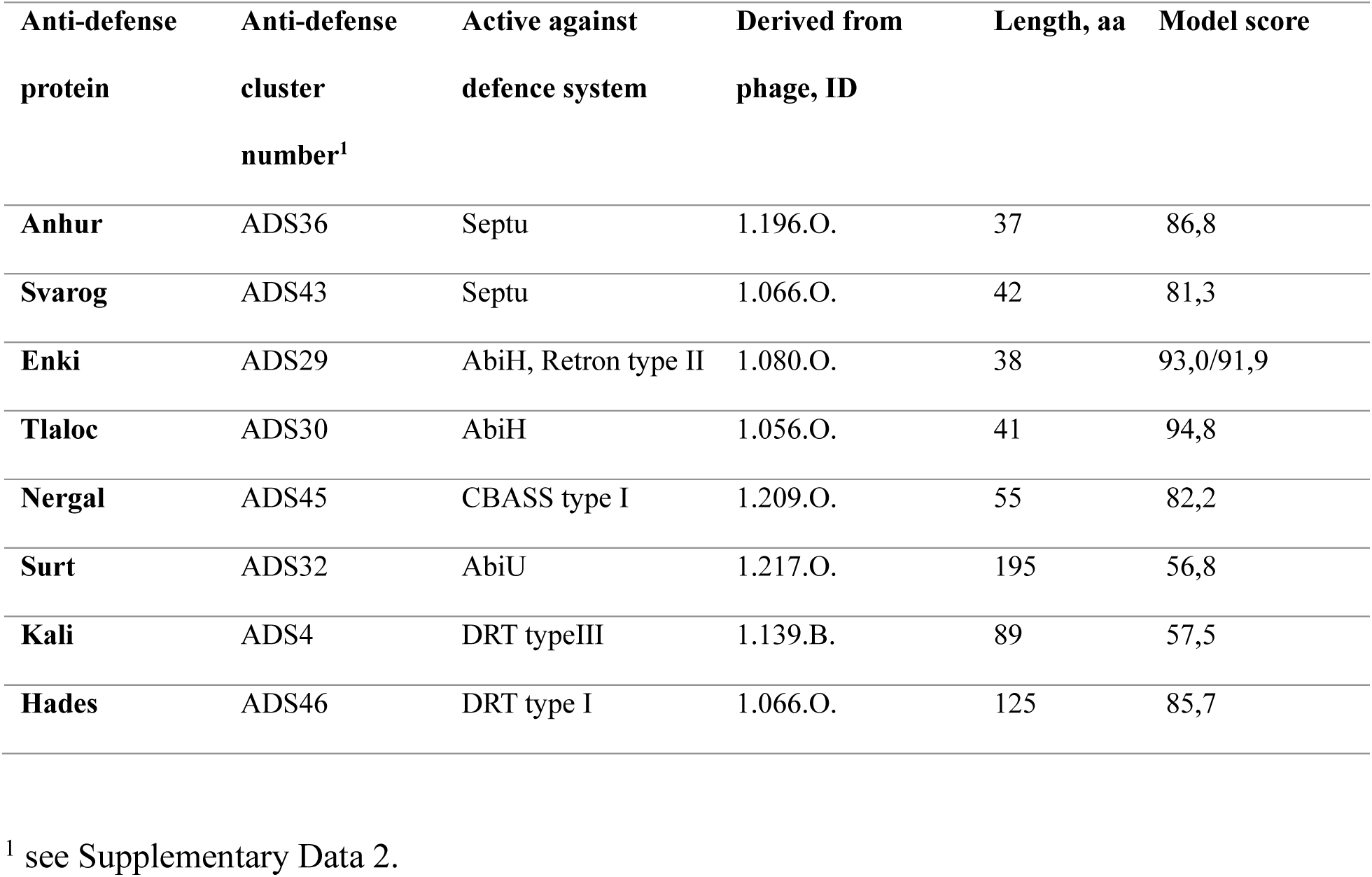
Experimentally verified anti-defense proteins active against a wide range of defense systems.

**Supplementary Table 6.** Examples of possible names for anti-defenses in the proposed naming scheme based on gods or mythological figures to whom some form of destruction is ascribed.

| **God<sup>1</sup>** | **Ascribed role in religion or mythology<sup>2</sup>** |
| --- | --- |
| **Shiva** | In Hindu mythology, Shiva is often depicted as the destroyer of the universe, responsible for bringing about the end of cycles of creation. |
| **Sekhmet** | In Egyptian mythology, Sekhmet is a lioness-headed goddess associated with war, destruction, and healing. |
| **Hades** | In Greek mythology, Hades is the god of the underworld, associated with death and the afterlife. |
| **Kali** | In Hindu mythology, Kali is a fierce goddess associated with destruction, time, and change. |
| **Thor** | In Norse mythology, Thor is the god of thunder associated with storms, lightning, and destruction. |
| **Anhur** | In Egyptian mythology, Anhur is a warrior god associated with war and destruction. |
| **Enki** | In Sumerian mythology, Enki is a god associated with water, wisdom, and also with the power to bring about floods and destruction. |
| **Tlaloc** | In Aztec mythology, Tlaloc is the god of rain, storms, and water, sometimes associated with destructive floods. |
| **Ares** | In Greek mythology, Ares is the god of war, violence, and bloodshed, often associated with destruction on the battlefield. |
| **Mars** | In Roman mythology, Mars is the counterpart of Ares, also representing war and destruction. |
| **Set** | In Egyptian mythology, Set is a god associated with chaos, storms, and destruction. He is often depicted as a malevolent force. |
| **Surt** | In Norse mythology, Surt is a fire giant who plays a significant role in the destruction of the world during Ragnarok, the end of the world. |
| **Eris** | In Greek mythology, Eris is the goddess of strife, discord, and chaos, often causing destruction through her actions. |
| *Nergal* | In Mesopotamian mythology, Nergal is the god of war and plague, associated with destruction and death. |
| *Svarog* | In Slavic mythology, Svarog is a deity associated with fire, blacksmithing, and sometimes destruction. |
| **Tezcatlipoca** | In Aztec mythology, Tezcatlipoca is a god associated with night, sorcery, and destruction, often depicted with a smoking mirror representing destruction and fate. |
| **Kraken** | A legendary sea monster known for its strength and ability to defeat sailors. |
| **Perseus** | A hero in Greek mythology who defeated the Gorgon Medusa with clever tactics. |
| **Fenrir** | A monstrous wolf in Norse mythology known for breaking free of chains and causing chaos. |
| **Cu Chulainn** | A hero in Irish mythology known for his exceptional skill in battle. |
| **Tiamat** | A primordial goddess in Babylonian mythology associated with chaos and overcoming order. |
| **Karna** | A warrior in Hindu mythology known for his skill and resilience in battle. |
| **Echidna** | A creature in Greek mythology known as the "Mother of Monsters," overcoming various foes. |

<sup>1</sup> Names used for systems verified here are italicized.
<sup>2</sup> The description was obtained from ChatGTP (March 2024)

**Supplementary table 7.**
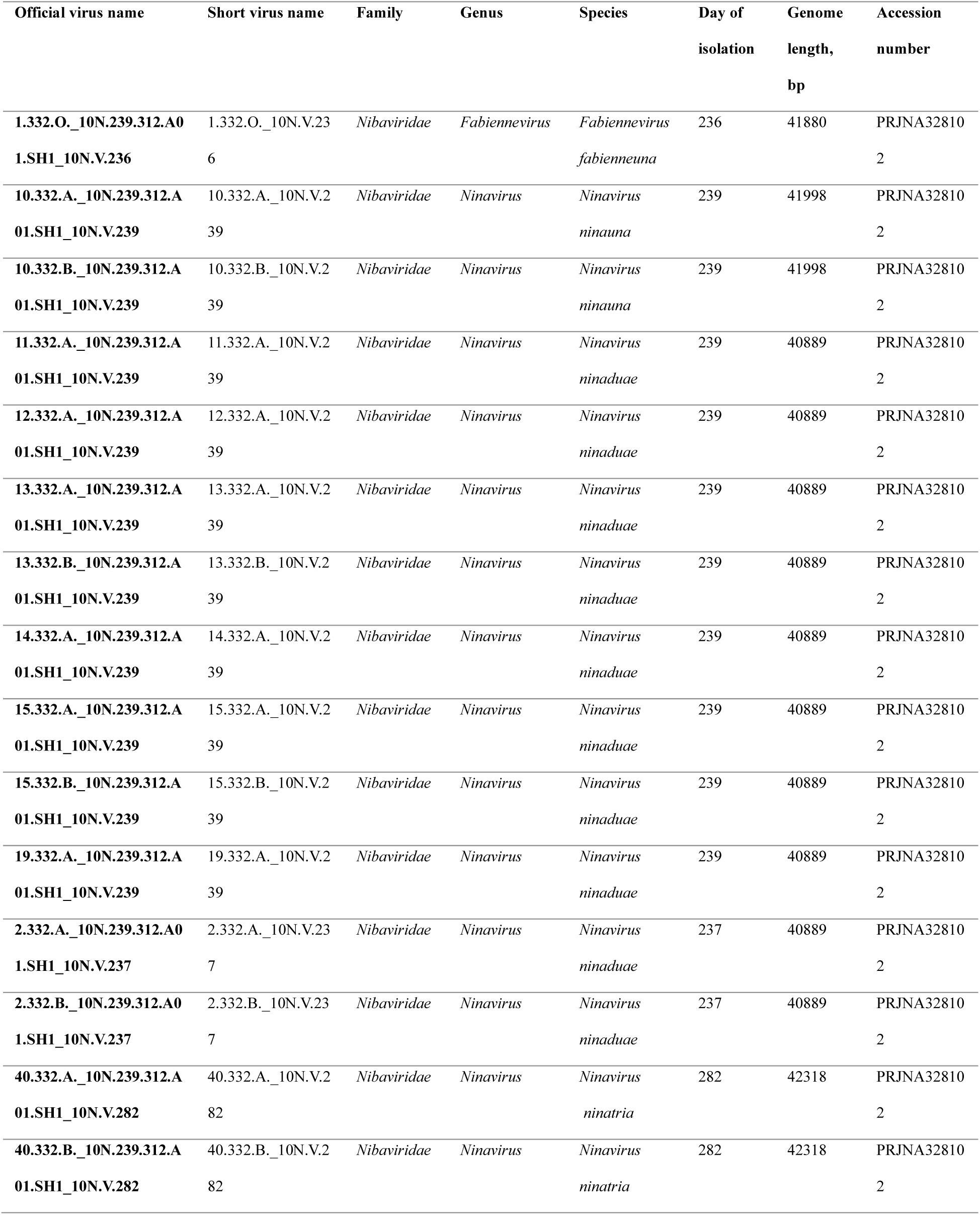
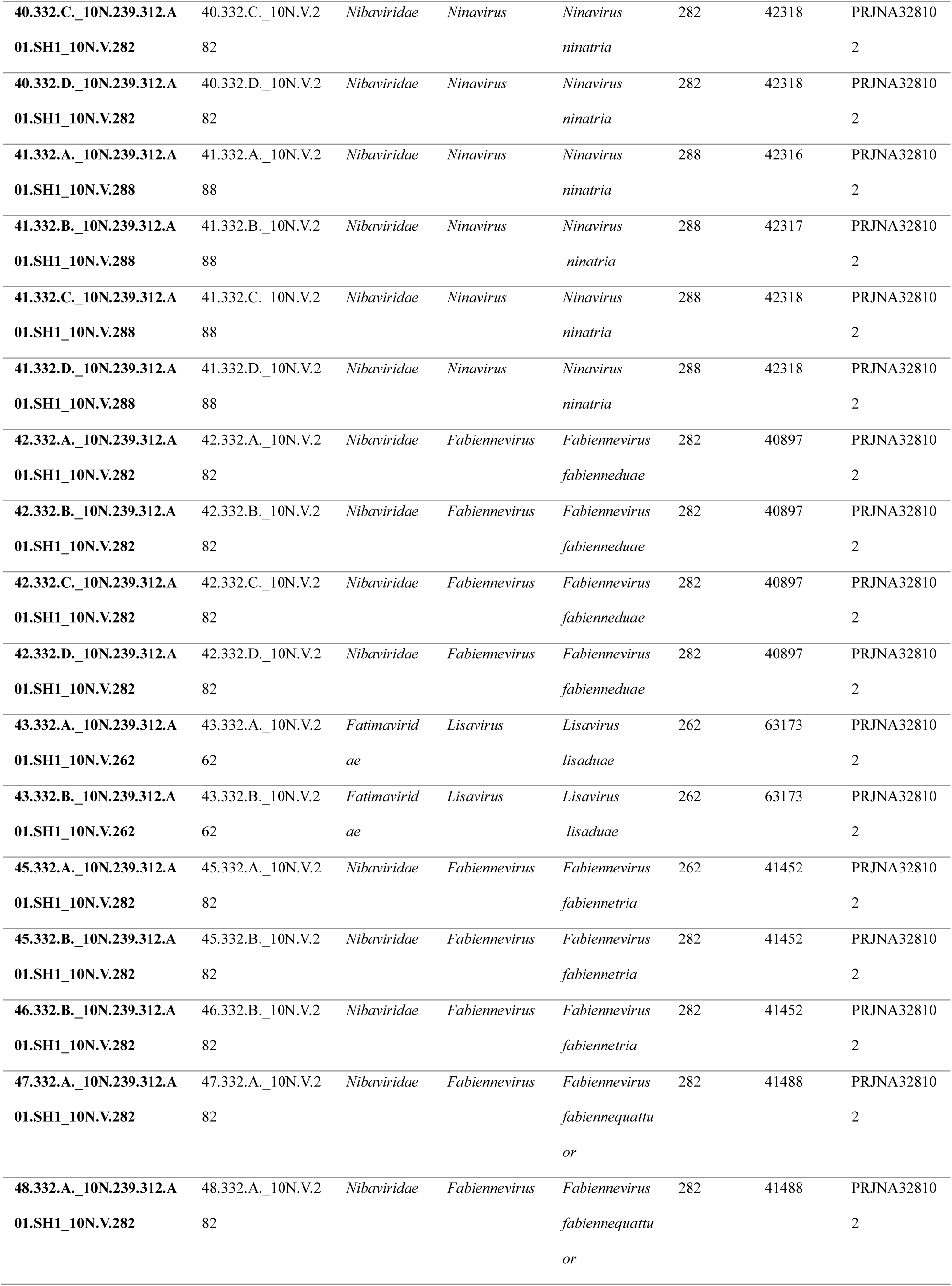
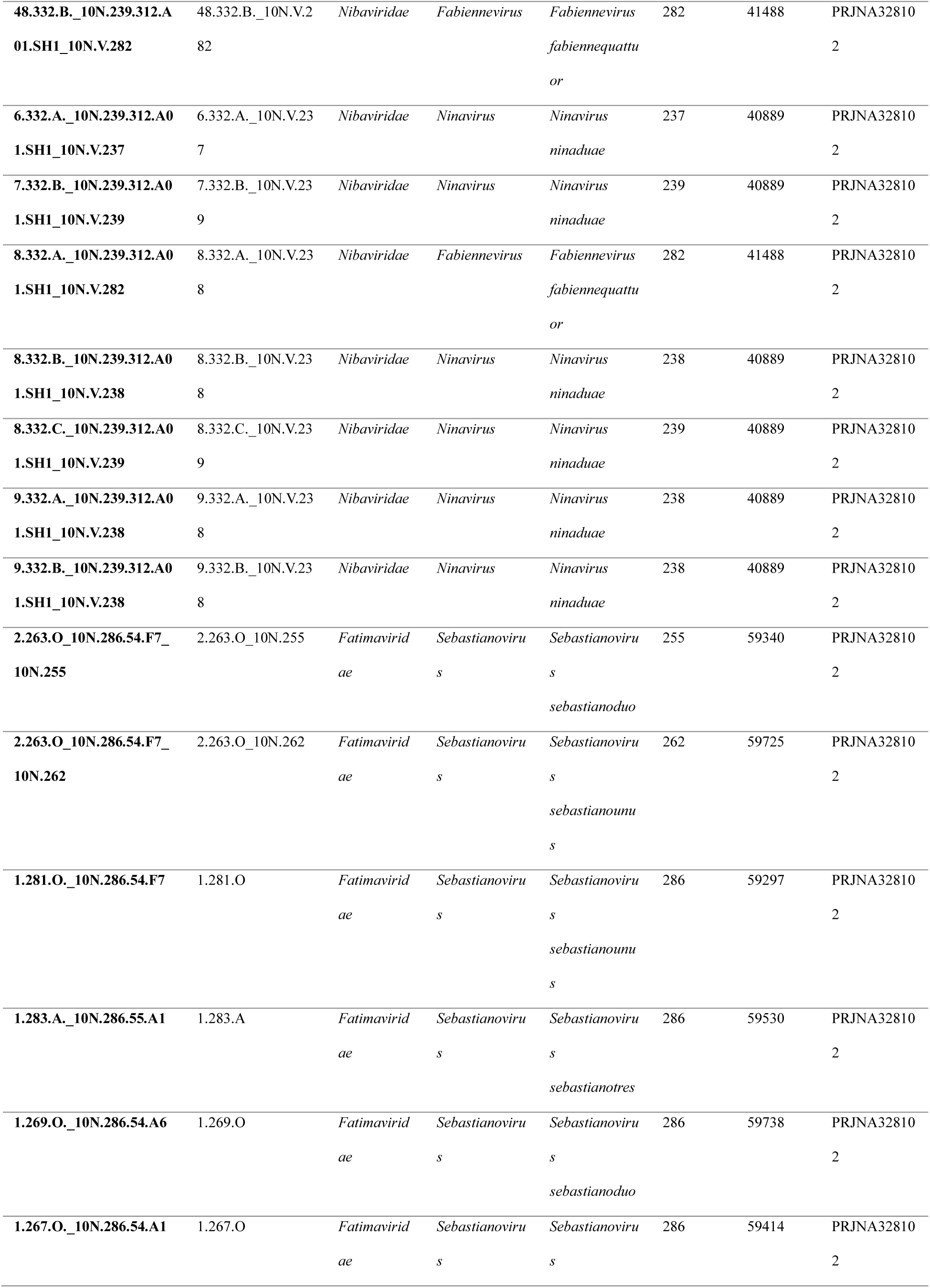
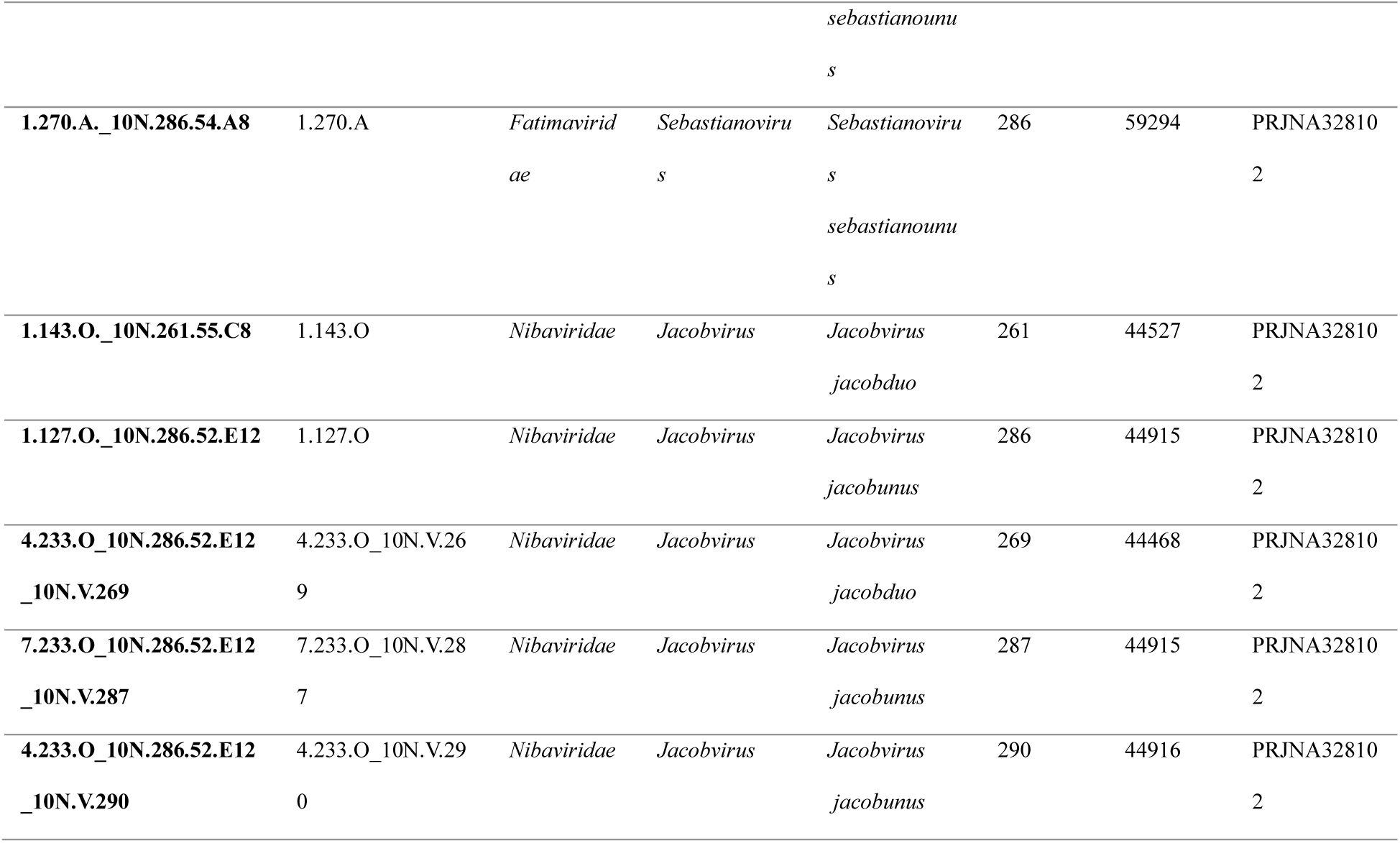
Classification of *V.cyclitrophicus* and *V.lentus* phages used in the plaque assays.

### Supplementary Data

**Supplementary Data 1.** Description of defense systems predicted in 758 Vibrio genomes using DefenseFInder v.1.0.8.

**Supplementary Data 2.** Putative anti-defense proteins predicted by the deep-learning model.

**Supplementary Data 3.** Results of LC-MS analysis.

**Supplementary Data 4.** Lists of bacterial and viral strains used in this study.

**Supplementary Data 5.** Annotation of viral genomes used in this study.

**Supplementary Data 6.** Annotation of phage defense elements (PDEs) knocked out of the genome of *Vibrio cyclitrophicus* 10N.239.312.A01

**Supplementary Data 7.** Detailed information on strains, plasmids and primers used in the study.

## Main References

1. Suttle, C. A. Marine viruses - Major players in the global ecosystem. Nat. Rev. Microbiol. 5, 801–812 (2007).

2. Breitbart, M. Marine viruses: Truth or dare. Annu. Rev. Mar. Sci. 4, 425–448 (2012).

3. Hussain, F. A. et al. Rapid evolutionary turnover of mobile genetic elements drives bacterial resistance to phages. Science 374, 488–492 (2021).

4. Tesson, F. et al. Systematic and quantitative view of the antiviral arsenal of prokaryotes. Nat. Commun. 13, (2022).

5. Georjon, H. & Bernheim, A. The highly diverse antiphage defense systems of bacteria. Nat. Rev. Microbiol. 21, 686–700 (2023).

6. Bernheim, A. & Sorek, R. The pan-immune system of bacteria: antiviral defense as a community resource. Nat. Rev. Microbiol. 18, 113–119 (2020).

7. Piel, D. et al. Genetic determinism of phage-bacteria coevolution in natural populations. bioRxiv 2021.05.05.442762 (2021) doi:10.1101/2021.05.05.442762.

8. Kauffman, K. M. et al. Resolving the structure of phage–bacteria interactions in the context of natural diversity. Nat. Commun. 13, 1–20 (2022).

9. Gaborieau, B. et al. Predicting phage-bacteria interactions at the strain level from genomes. bioRxiv 2023.11.22.567924 (2023).

10. Costa, A. R. et al. Accumulation of defense systems in phage-resistant strains of Pseudomonas aeruginosa. Sci. Adv. 10, eadj0341 (2024).

11. Pinilla-Redondo, R. et al. Discovery of multiple anti-CRISPRs highlights anti-defense gene clustering in mobile genetic elements. Nat. Commun. 11, 5652 (2020).

12. Piel, D. et al. Phage–host coevolution in natural populations. Nat. Microbiol. 2022 7, 1– 12 (2022).

13. Mayo-Muñoz, D., Pinilla-Redondo, R., Camara-Wilpert, S., Birkholz, N. & Fineran, P. C. Inhibitors of bacterial immune systems: discovery, mechanisms and applications. Nat. Rev. Genet. (2024) doi:10.1038/s41576-023-00676-9.

14. Sukrit Silas, Héloïse Carion, Kira S. Makarova, Eric Laderman, David Sanchez Godinez, Matthew Johnson, Andrea Fossati Danielle Swaney, Michael Bocek, Eugene V.Koonin, J. B.-D. Activation of programmed cell death and counter-defense functions of phage accessory genes. (2023).

15. Huiting, E. et al. Bacteriophages inhibit and evade cGAS-like immune function in bacteria. Cell 186, 864–876.e21 (2023).

16. Hobbs, S. J. et al. Phage anti-CBASS and anti-Pycsar nucleases subvert bacterial immunity. Nature 605, 522–526 (2022).

17. Jenson, J. M., Li, T., Du, F., Ea, C. K. & Chen, Z. J. Ubiquitin-like conjugation by bacterial cGAS enhances anti-phage defense. Nature 616, 326–331 (2023).

18. Leavitt, A. et al. Viruses inhibit TIR gcADPR signalling to overcome bacterial defense. Nature 611, 326–331 (2022).

19. Yirmiya, E. et al. Phages overcome bacterial immunity via diverse anti-defense proteins. Nature 0–1 (2023) doi:10.1038/s41586-023-06869-w.

20. Garb, J. et al. Multiple phage resistance systems inhibit infection via SIR2-dependent NAD+ depletion. Nat. Microbiol. 7, 1849–1856 (2022).

21. Antine, S. P. et al. Structural basis of Gabija anti-phage defense and viral immune evasion. Nature (2023) doi:10.1038/s41586-023-06855-2.

22. Bepler, T. & Berger, B. Learning the protein language: Evolution, structure, and function. Cell Syst. 12, 654–669.e3 (2021).

23. Chowdhury, R. et al. Single-sequence protein structure prediction using a language model and deep learning. Nat. Biotechnol. 40, 1617–1623 (2022).

24. Zhang, Z. et al. Protein language models learn evolutionary statistics of interacting sequence motifs. 2024.01.30.577970 Preprint at 10.1101/2024.01.30.577970 (2024).

25. Devlin, J., Chang, M.-W., Lee, K. & Toutanova, K. BERT: Pre-training of Deep Bidirectional Transformers for Language Understanding. Preprint at 10.48550/arXiv.1810.04805 (2019).

26. Lundberg, S. M. & Lee, S.-I. A Unified Approach to Interpreting Model Predictions. in Advances in Neural Information Processing Systems vol. 30 (Curran Associates, Inc., 2017).

27. Bobonis, J. et al. Bacterial retrons encode phage-defending tripartite toxin–antitoxin systems. Nature 609, 144–150 (2022).

28. Doron, S. et al. Systematic discovery of antiphage defense systems in the microbial pangenome. Science eaar4120 (2018) doi:10.1126/science.aar4120.

29. Prévots, F., Daloyau, M., Bonin, O., Dumont, X. & Tolou, S. Cloning and sequencing of the novel abortive infection gene abiH of Lactococcus lactis ssp. lactis biovar. diacetylactis S94. FEMS Microbiol. Lett. 142, 295–299 (1996).

30. Zaremba, M. et al. Short prokaryotic Argonautes provide defense against incoming mobile genetic elements through NAD+ depletion. Nat. Microbiol. 7, 1857–1869 (2022).

31. Makarova, K. S., Wolf, Y. I., van der Oost, J. & Koonin, E. V. Prokaryotic homologs of Argonaute proteins are predicted to function as key components of a novel system of defense against mobile genetic elements. Biol. Direct 4, 29 (2009).

32. Ofir, G. et al. Antiviral activity of bacterial TIR domains via immune signalling molecules. Nature 600, 116–120 (2021).

33. Gao, L. et al. Diverse Enzymatic Activities Mediate Antiviral Immunity in Prokaryotes. Science 369, 1077 (2020).

## Methods References

34. Schwengers, O. et al. Bakta: rapid and standardized annotation of bacterial genomes via alignment-free sequence identification. Microb. Genomics 7, 000685 (2021).

35. McNair, K., Zhou, C., Dinsdale, E. A., Souza, B. & Edwards, R. A. PHANOTATE: a novel approach to gene identification in phage genomes. Bioinformatics 35, 4537–4542 (2019).

36. Moraru, C. VirClust—A Tool for Hierarchical Clustering, Core Protein Detection and Annotation of (Prokaryotic) Viruses. Viruses 15, 1007 (2023).

37. Zayed, A. A. et al. efam: an expanded, metaproteome-supported HMM profile database of viral protein families. Bioinformatics 37, 4202–4208 (2021).

38. Terzian, P., et al. PHROG: families of prokaryotic virus proteins clustered using remote homology. NAR Genomics Bioinforma. 3, lqab067 (2021).

39. Kiening, M. et al. Conserved Secondary Structures in Viral mRNAs. Viruses 11, 401 (2019).

40. Grazziotin, A. L., Koonin, E. V. & Kristensen, D. M. Prokaryotic Virus Orthologous Groups (pVOGs): a resource for comparative genomics and protein family annotation. Nucleic Acids Res. 45, D491–D498 (2017).

41. Jones, P. et al. InterProScan 5: genome-scale protein function classification. Bioinformatics 30, 1236–1240 (2014).

42. Chan, P. P., Lin, B. Y., Mak, A. J. & Lowe, T. M. tRNAscan-SE 2.0: improved detection and functional classification of transfer RNA genes. Nucleic Acids Res. 49, 9077–9096 (2021).

43. Laslett, D. & Canback, B. ARAGORN, a program to detect tRNA genes and tmRNA genes in nucleotide sequences. Nucleic Acids Res. 32, 11–16 (2004).

44. Brettin, T. et al. RASTtk: a modular and extensible implementation of the RAST algorithm for building custom annotation pipelines and annotating batches of genomes. Sci. Rep. 5, 8365 (2015).

45. Altschul, S. F., Gish, W., Miller, W., Myers, E. W. & Lipman, D. J. Basic local alignment search tool. J. Mol. Biol. 215, 403–410 (1990).

46. Cook, R. et al. INfrastructure for a PHAge REference Database: Identification of Large-Scale Biases in the Current Collection of Cultured Phage Genomes. PHAGE New Rochelle N 2, 214–223 (2021).

47. Nishimura, Y. et al. ViPTree: the viral proteomic tree server. Bioinforma. Oxf. Engl. 33, 2379–2380 (2017).

48. Moraru, C., Varsani, A. & Kropinski, A. M. VIRIDIC—A Novel Tool to Calculate the Intergenomic Similarities of Prokaryote-Infecting Viruses. Viruses 12, 1268 (2020).

49. Traag, V. A., Waltman, L. & van Eck, N. J. From Louvain to Leiden: guaranteeing well-connected communities. Sci. Rep. 9, 5233 (2019).

50. Buchfink, B., Reuter, K. & Drost, H.-G. Sensitive protein alignments at tree-of-life scale using DIAMOND. Nat. Methods 18, 366–368 (2021).

51. Paszke, A. et al. PyTorch: an imperative style, high-performance deep learning library. in Proceedings of the 33rd International Conference on Neural Information Processing Systems 8026–8037 (Curran Associates Inc., Red Hook, NY, USA, 2019).

52. Chaos Game Representation | SIAM Review. https://epubs.siam.org/doi/10.1137/20M1386438.

53. R Core Team. R: A Language and Environment for Statistical Computing. R Foundation for Statistical Computing, Vienna, Austria. https://www.r-project.org/. (2020).

54. Kauffman, K. M. et al. Viruses of the Nahant Collection, characterization of 251 marine Vibrionaceae viruses. Sci. Data 5, 180114 (2018).

55. Martin-Platero, A. M. et al. High resolution time series reveals cohesive but short-lived communities in coastal plankton. Nat. Commun. 9, 266 (2018).

56. Kauffman, K. M. et al. A major lineage of non-tailed dsDNA viruses as unrecognized killers of marine bacteria. Nature 554, 118–122 (2018).

57. Hunt, D. E. et al. Resource partitioning and sympatric differentiation among closely related bacterioplankton. Science 320, 1081–1085 (2008).

58. Kolmogorov, M., Yuan, J., Lin, Y. & Pevzner, P. A. Assembly of long, error-prone reads using repeat graphs. Nat. Biotechnol. 37, 540–546 (2019).

59. Gurevich, A., Saveliev, V., Vyahhi, N. & Tesler, G. QUAST: quality assessment tool for genome assemblies. Bioinforma. Oxf. Engl. 29, 1072–1075 (2013).

60. Parks, D. H., Imelfort, M., Skennerton, C. T., Hugenholtz, P. & Tyson, G. W. CheckM: assessing the quality of microbial genomes recovered from isolates, single cells, and metagenomes. Genome Res. 25, 1043–1055 (2015).

61. Gautreau, G. et al. PPanGGOLiN: Depicting microbial diversity via a partitioned pangenome graph. PLOS Comput. Biol. 16, e1007732 (2020).

62. Arndt, D. et al. PHASTER: a better, faster version of the PHAST phage search tool. Nucleic Acids Res. 44, W16–W21 (2016).

63. Guo, J. et al. VirSorter2: a multi-classifier, expert-guided approach to detect diverse DNA and RNA viruses. Microbiome 9, 37 (2021).

64. Néron, B. et al. MacSyFinder v2: Improved modelling and search engine to identify molecular systems in genomes. Peer Community J. 3, (2023).

65. Tesson, F. et al. A Comprehensive Resource for Exploring Antiphage Defense: DefenseFinder Webservice, Wiki and Databases. 2024.01.25.577194 Preprint at 10.1101/2024.01.25.577194 (2024).

66. Bobonis, J., Yang, A. L. J., Voogdt, C. G. P. & Typas, A. TAC–TIC, a high-throughput genetics method to identify triggers or blockers of bacterial toxin–antitoxin systems. Nat. Protoc. 1–19 (2024) doi:10.1038/s41596-024-00988-y.

67. Le Roux, F., Davis, B. M. & Waldor, M. K. Conserved small RNAs govern replication and incompatibility of a diverse new plasmid family from marine bacteria. Nucleic Acids Res. 39, 1004–1013 (2011).

68. Lefort, V., Desper, R. & Gascuel, O. FastME 2.0: A Comprehensive, Accurate, and Fast Distance-Based Phylogeny Inference Program. Mol. Biol. Evol. 32, 2798–2800 (2015).

69. Wickham, H. Ggplot2. (Springer International Publishing, Cham, 2016). doi:10.1007/978-3-319-24277-4.

